# LDB1-dependent enhancer connectivity defines T-cell leukemia identities and masks metabolic vulnerabilities

**DOI:** 10.64898/2026.06.29.733236

**Authors:** Rahul S Bhansali, Juan Sebastian Long, Petri Pölönen, Tomoya Isobe, Siqing Wang, Shuo Zhang, Zhuangzhuang Geng, Ahnaf Tausif, Mariah C Antopia, Nicholas G Aboreden, Sarah J Skuli, Belinda M Giardine, Cheryl A Keller, Ross C Hardison, Kai Tan, Charles G Mullighan, Gerd A Blobel

**Author notes:** Corresponding Author(s): Rahul S Bhansali, MD Abramson Research Center 315 3615 Civic Center Blvd. Philadelphia, PA 19104-4318, Gerd A Blobel, MD, PhD Abramson Research Center 316H 3615 Civic Center Blvd., Philadelphia, PA 19104-4318.

## Abstract

Spatial enhancer connectivity is fundamental to proper gene regulation. Enhancer dysregulation has emerged as a hallmark of cancers, including T-cell acute lymphoblastic leukemias (T-ALL). T-ALL are aggressive malignancies characterized by marked transcriptional heterogeneity driven by distinct stages of developmental arrest and diverse noncoding alterations. How these cancers co-opt nuclear architecture to rewire enhancer connectivity remains poorly understood. Here, we report that the LDB1 chromatin architectural complex is an essential mediator of enhancer-oncogene looping that sustains oncogenic transcriptional programs across multiple T-ALL subtypes. Integrating bulk and single-cell transcriptomic data from patients with T-ALL and healthy hematopoietic controls, we show that the LDB1-dependent regulatory circuitry defines the molecular identities of distinct T-ALL subtypes while restricting plasticity toward alternative cell states. LDB1 loss dismantles chromatin looping among cell state-defining enhancers liberating them to form promiscuous interactions with nearby genes. This enhancer rewiring stimulates expression of key metabolic genes, creating a mevalonate pathway dependency exploitable with statin treatment. Our study establishes LDB1 as a central executor of T-ALL regulatory circuitry and more broadly illustrates chromatin rewiring as a source of targetable dependencies in cancer.

## Introduction

The 3D organization of the mammalian genome impacts and is impacted by all nuclear processes, including gene transcription, DNA replication and repair (reviewed in ^1–3^). Dysregulation of factors that sculpt the genome can cause developmental diseases and cancer. Central to the establishment of cell type-specific gene expression patterns is the proper spatial connectivity between cis-regulatory elements (CREs) such as enhancers and gene promoters^4^. While universal mechanisms involving the cohesin machinery and CTCF contribute to genome folding, CRE connectivity requires additional architectural factors that are employed in a gene-specific fashion (reviewed in ^5^). Gain- or loss-of-function of such factors can lead to enhancer-promoter miswiring, triggering widespread changes in gene transcription, potentially causing malignant transformation. Prominent examples include Lim-domain only 2 (LMO2) and its paralog LMO1, which were among the earliest recurrent chromosomal rearrangements identified in T-cell acute lymphoblastic leukemia (T-ALL)^6–9^. LMO2 is an essential component of a molecular assembly that includes the architectural factor LDB1, which can dimerize or oligomerize to promote chromatin loops among LDB1/LMO2-occupied regulatory elements^10–29^. LDB1 complexes are in turn stabilized by SSBP (single stranded DNA binding protein) family proteins. Neither LDB1 nor SSBP proteins bind DNA directly but are rather tethered to DNA via sequence-specific transcription factors (TFs) with LMO2 serving as a bridge (**Fig. 1a**).

**Figure 1.**
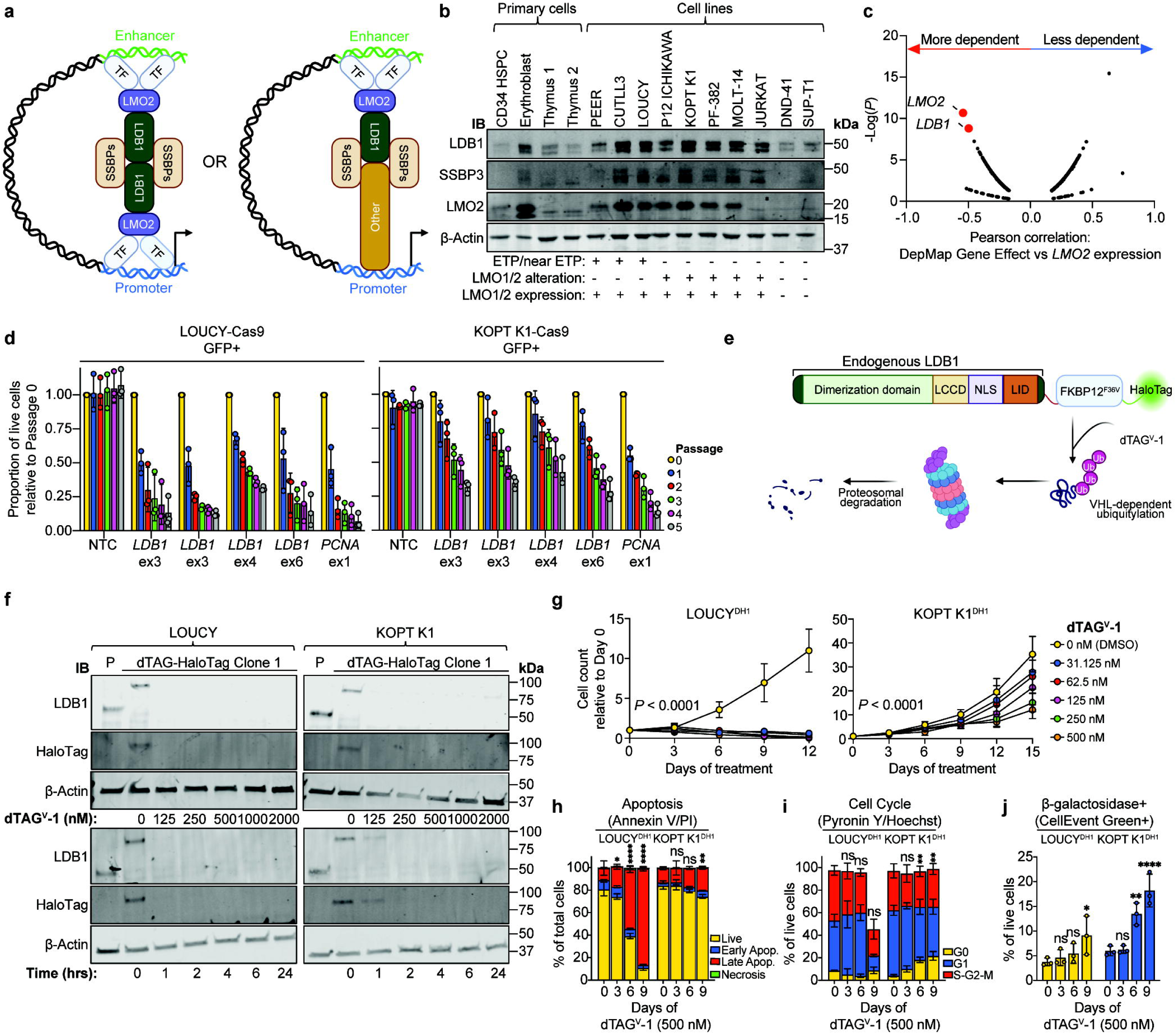
LDB1 is a dependency in *LMO1/2*-expressing T-ALL. **a,** Schematic of LDB1-dependent chromatin looping. **b,** Representative Western blot depicts LDB1, SSBP3, LMO2 and β-Actin protein expression in human primary hematopoietic tissues and human T-ALL cell lines. **c,** Pearson correlation plot for DepMap Chronos Gene Effect Score (dependency strength) versus *LMO2* expression. **d,** Proportion of GFP+ LOUCY- or KOPT K1-Cas9 cells after transduction with non-targeting control (NTC), *PCNA*-targeting (positive control) or *LDB1*-targeting sgRNAs; values normalized to Passage 0 for each sgRNA. **e,** Schematic depicting LDB1-dTAG-HaloTag construct and degradation mechanism. **f,** Representative Western blot depicts LDB1, HaloTag and β-Actin protein expression in LOUCY and KOPT K1 parental cells (P) and dTAG-HaloTag subclones; top, dTAG^V^-1 dose titration (4-hour treatment); bottom, dTAG^V^-1 time titration (500 nM). **g,** LOUCY^DH1^ and KOPT K1^DH1^ cell counts upon serial dTAG^V^-1 treatment (0-500 nM); cell counts normalized to Day 0; *P*-values from two-way ANOVA with post hoc Tukey test. **h,** Distribution of live, early apoptotic, late apoptotic and necrotic cells upon serial dTAG^V^-1 treatment, defined by Annexin V/Propidium Iodide (PI) (gating in **Supplemental Fig. S2a**); *P*-values from one-way ANOVA with post hoc Dunnett’s test comparing late apoptotic populations to Day 0. **i,** Distribution of G0, G1 and S-G2-M cells upon serial dTAG^V^-1 treatment, defined by Pyronin Y/Hoechst 33342 (gating in **Supplemental Fig. S2b**); *P*-values from one-way ANOVA with post hoc Dunnett’s test comparing G0 populations to Day 0. **j,** Percent of β-galactosidase+ cells upon serial dTAG^V^-1 treatment, defined by CellEvent Senescence Green (gating in **Supplemental Fig. S2c**); *P*-values from one-way ANOVA with post hoc Dunnett’s test relative to Day 0. *N =* 3 biological replicates for all experiments; for primary thymic samples prepared from cryopreserved tissue, replicate aliquots were prepared from the same tissue extracts. TF, transcription factor; HSPC, hematopoietic stem and progenitor cell; ETP, early thymic precursor; IB, immunoblot; ex, Exon; LCCD, LDB1/Chip conserved domain; NLS, nuclear localization sequence; LID, Lim-interacting domain; Ub, Ubiquitination.

In normal development, LDB1/LMO2-nucleated complexes are central to the formation of enhancer-promoter loops at hematopoietic gene loci and are essential for hematopoietic stem cell development^12–14,16–23^. Although LDB1 and LMO2 are dispensable for normal thymopoiesis, forced *Lmo2* overexpression in mice results in a highly penetrant and LDB1-dependent T-ALL^30–33^. Furthermore, diverse noncoding alterations^34–39^ drive *LMO2* overexpression in over 50% of T-ALL cases across multiple differentiation states and molecular subtypes, together implicating a broader role for LDB1/LMO2 in T-ALL pathogenesis than previously appreciated.

T-ALL accounts for approximately 15% and 25% of pediatric and adult cases of ALL, respectively^40^. While outcomes have improved with contemporary treatment regimens, relapsed/refractory disease is common and associated with a dismal prognosis^41,42^. *NOTCH1* and *CDKN2A* alterations are common across most T-ALL subtypes^43,44^, impacting around 70% of cases. However, a recent comprehensive multi-omic analysis^36^ found a comparable prevalence of noncoding alterations resulting in enhancer-mediated activation of over 60 oncogenes, underscoring the importance of enhancer dysregulation in T-ALL pathogenesis^45–47^. This raises key questions about the basis for LDB1/LMO2 as leukemic drivers across T-ALL subtypes. Specifically, how does LDB1/LMO2-mediated spatial clustering among CREs support oncogenic transcriptional programs, and does such CRE clustering constrain regulatory elements from activating alternative programs that may represent new therapeutically actionable pathways?

To address these questions, we devised an LDB1 degron system in human T-ALL cell lines representing distinct T-ALL subtypes coupled with analyses in primary human T-ALL datasets to define LDB1-dependent oncogenic enhancer connectivity and transcriptional programs. We show that LDB1 spatially connects regulatory elements that define T-ALL subtype features beyond merely preventing T-cell maturation. By physically clustering enhancers, LDB1 restricts their promiscuous rewiring and associated alternative gene expression programs. Based on this paradigm, we propose that enhancer rewiring may serve as a source of new therapeutic vulnerabilities in cancer.

## Results

### LDB1 is a dependency in T-cell acute lymphoblastic leukemia with LMO1/2 expression

*Lmo2* is highly expressed in murine hematopoietic stem and progenitor cells (HSPCs) but during normal thymopoiesis is silenced after the early thymic precursor (ETP) stage, coinciding with definitive commitment to the T-cell lineage^48^. This developmental checkpoint is critical because historically, T-ALL have been classified as ETP/near-ETP or non-ETP, with the former harboring high-risk HSPC-like features and often lacking canonical T-ALL driver alterations such as *NOTCH1* mutations^36,49^. We analyzed RNA-seq data from human postnatal thymocytes^50^ and observed the expected decline of *LMO2* expression during thymopoiesis, whereas *LDB1* expression is retained (**Supplemental Fig. S1a**). We next characterized LMO2 and LDB1 protein levels in human T-ALL cell lines (**Supplemental Table 1**) as well as normal hematopoietic counterparts (CD34+ HSPCs, proerythroblast cells, pediatric whole thymic tissue). LDB1 and its cofactor SSBP3^22,51^ are elevated in T-ALL subtypes with ETP/near-ETP phenotypes as well as non-ETP subtypes with LMO1/2-activating alterations, when compared with LMO1/2-deficient lines and normal hematopoietic controls (**Fig. 1b**). This predicts that LMO2 overexpression is capable of forming functional architectural complexes in T-ALL.

To examine *LMO2* expression as a cancer dependency, we analyzed all hematologic cancer cell lines in the DepMap Portal^52^ (Broad Institute) and confirmed LDB1 as a bona fide co-dependency (**Fig. 1c**). However, since few T-ALL cell lines are represented in the DepMap CRISPR knockout pipeline, we used CRISPR/Cas9 to knockout LDB1 in 5 T-ALL cell lines. We found that LDB1 dependency was strongest in those with LMO1/2 expression, with LDB1 knockout leading to progressive loss of T-ALL cell expansion in all cell lines except SUP-T1, which is LMO1/2-deficient (**Fig. 1d**; **Supplemental Fig. S1b, c**).

We previously demonstrated that LDB1 is required for maintenance of a considerable subset of CRE loops in a manner not involving cohesin and CTCF^21^. Moreover, forced tethering of LDB1 to chromatin is sufficient to induce chromatin loop formation^18–22^. Given the broad and essential role of LDB1 in connecting CREs, we set out to define which architectural elements and transcriptional targets are regulated by the LDB1/LMO2 complex. We engineered isogenic T-ALL cell lines with FKBP12^F36V^ (dTAG) and HaloTag homozygously fused in tandem to the endogenous LDB1 C-terminus (**Fig. 1e**). The LOUCY and KOPT K1 cell lines were selected as model systems, as they reflect distinct differentiation states (ETP versus near late cortical, respectively), molecular drivers (*SET*::*NUP214* versus *TRA/TRD::LMO2*, respectively) and mechanisms of LMO2 activation (developmental arrest versus enhancer hijacking, respectively), together providing distinct cellular contexts of LDB1 and LMO2 cooperativity in T-cell leukemogenesis. Since insertion of a bulky tag may impact the function of a transcriptional coregulator, we selected two clones each, performed RNA-seq and found minimal differences in transcriptomes between LOUCY-dTAG-HaloTag (LOUCY^DH^) and KOPT K1-dTAG-HaloTag (KOPT K1^DH^) subclones and their respective parental cell lines (**Supplemental Fig. S1d, e**), including key oncogenic driver genes (**Supplemental Fig. S1f**). Furthermore, treatment with dTAG^V^-1 ligand^53^, which targets dTAG-fused proteins to the von Hippel-Lindau (VHL) E3 ligase complex, depleted LDB1 protein by over 90% within less than 4 hours as determined by Western blot and flow cytometry using the J646 Janelia Fluor HaloTag Ligand (**Fig. 1f; Supplemental Fig. S1g**).

Prolonged LDB1 depletion impaired expansion of both cell lines in a dose-dependent manner (**Fig. 1g; Supplemental Fig. S1h**) with more potent anti-leukemic effects in LOUCY^DH^ cells compared with KOPT K1^DH^ cells, consistent with our LDB1 knockout experiments in parental cell lines. In LOUCY^DH^ cells, LDB1 loss induced apoptosis (**Fig. 1h; Supplemental Fig. S2a**) while in KOPT K1^DH^ cells, it elicited a cytostatic effect with cumulative G0 arrest (**Fig. 1i; Supplemental Fig. S2b**), increased β-galactosidase staining (**Fig. 1j; Supplemental Fig. S2c**), and upregulation of senescence-associated transcripts (**Supplemental Fig. S1i)**, together suggestive of a senescence-like phenotype.

Collectively, these findings underscore LDB1 as a critical dependency in T-ALL with LMO1/2 activation, though the mechanism of this dependency may vary between T-ALL subtypes.

### LDB1 occupies T-ALL subtype-specific cis-regulatory networks

To examine the basis for subtype-specific effects of LDB1 depletion on T-ALL growth arrest, we performed LDB1 ChIP-seq in LOUCY^DH^ cells and KOPT K1^DH^ cells. Indeed, LDB1 occupancy profiles significantly differed between cell lines with only 38% of peaks in LOUCY^DH^ (14,657/38,550) and 56.1% of peaks in KOPT K1^DH^ (14,428/25,706) overlapping with the other cell line (**Fig. 2a**). We also carried out ChIP-seq for LMO2 and SSBP3 in both cell lines and found that most peaks overlapped with LDB1 peaks, reflecting assembly of the canonical LDB1/LMO2 architectural complex (**Supplemental Fig. S3a, b**). Consistent with our prior work^21,22^, LDB1 is predominantly recruited to CREs, the majority being active enhancers, defined by H3K27ac peaks further than 1 kb from a transcription start site (TSS) (**Fig. 2b, c**). At enhancer regions, LDB1 occupancy showed an even greater divergence between cell lines, with only 30.9% of peaks in LOUCY^DH^ (4,798/15,535) and 41.9% of peaks in KOPT K1^DH^ (4,731/11,296) overlapping with the other cell line (**Fig. 2d, e**). Shared LDB1 peaks at enhancers were enriched for RUNX and ETS family motifs as expected^12,13^, while T-ALL subtype-specific LDB1 peaks at enhancers showed distinct motif patterns reflecting early progenitor identity in LOUCY^DH^ (GATA and HOX families) and T-cell signaling programs in KOPT K1^DH^ (STAT family and HEB) (**Supplemental Fig. S3c; Supplemental Table 2**).

**Figure 2.**
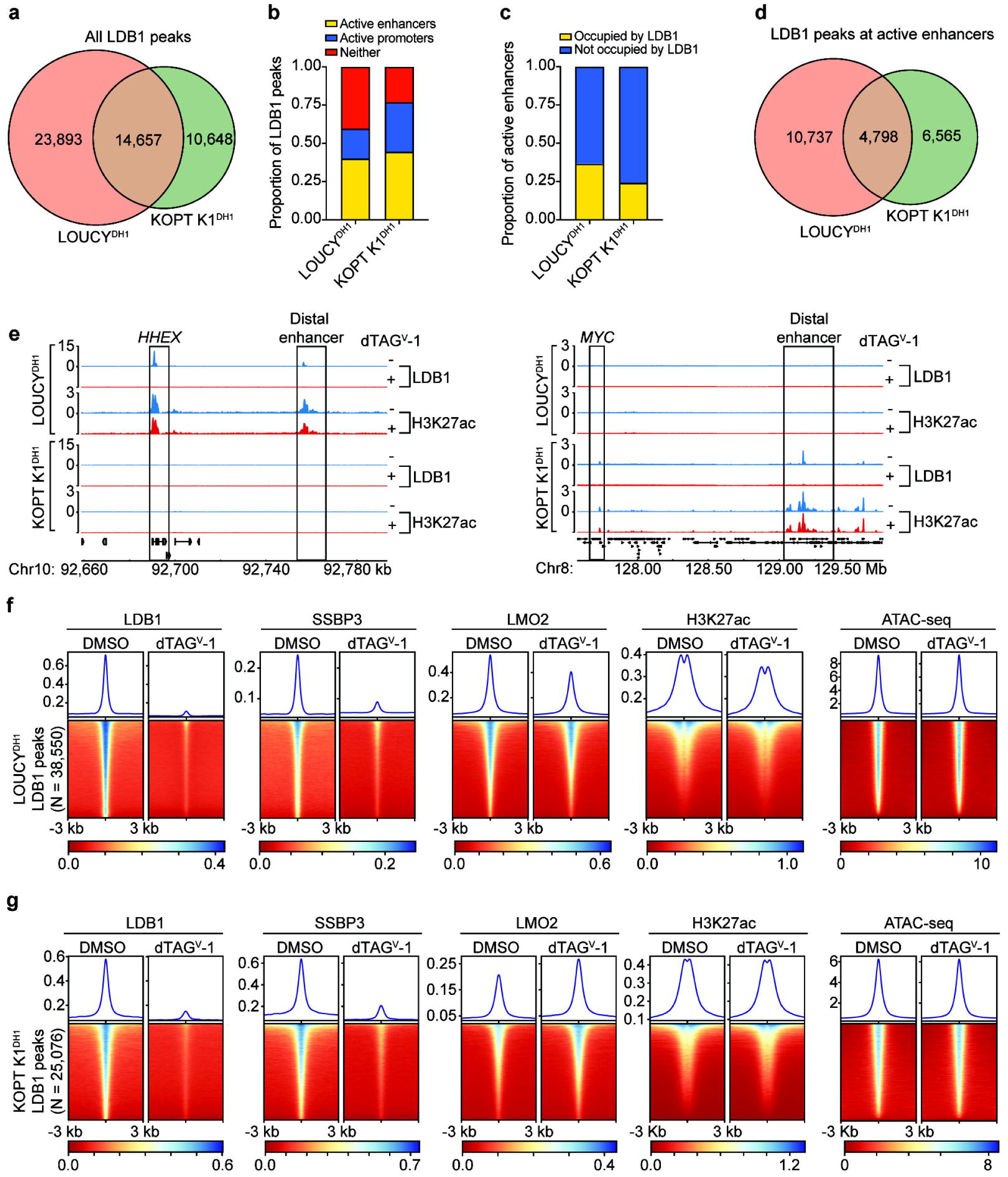
LDB1 recruitment to enhancers is specific to T-ALL subtypes. **a,** Overlap between LDB1 ChIP-seq peaks in LOUCY^DH1^ and KOPT K1^DH1^. **b,** Proportion of LDB1 ChIP-seq peaks overlapping with active enhancers, active promoters or neither. **c,** Proportion of active enhancers occupied by LDB1 ChIP-seq peaks. **d,** Overlap between LDB1 ChIP-seq peaks at active enhancers in LOUCY^DH1^ and KOPT K1^DH1^. **e,** Representative LDB1 and H3K27ac ChIP-seq tracks for LOUCY-specific and KOPT K1-specific LDB1-occupied enhancers (cyan, DMSO; red, dTAG^V^-1); **f,** Heatmaps of LOUCY^DH1^ ChIP-seq and ATAC-seq signal upon acute LDB1 depletion; profiles centered over all LDB1 peaks from DMSO-treated cells. **g**, Similar to **f,** but depicting KOPT K1^DH1^ ChIP-seq and ATAC-seq data. Biological replicates per cell line and treatment: LDB1 ChIP-seq, *N =* 4; LMO2 ChIP-seq, *N =* 2; SSBP3 ChIP-seq, *N =* 2; H3K27ac ChIP-seq, *N =* 2; ATAC-seq, *N =* 3.

In contrast to our findings in murine erythroid cells^21^, a larger fraction of LDB1 peaks at enhancers co-localized with cohesin (SMC3) in the absence of CTCF (52.2% in LOUCY^DH^ and 58.2% in KOPT K1^DH^) (**Supplemental Fig. S3d, e**), characteristic of highly active enhancer elements rather than topologically associating domain (TAD) boundaries which are typically co-occupied by CTCF and SMC3. Importantly, acute LDB1 depletion had minimal impact on H3K27ac signal, chromatin accessibility (as measured by ATAC-seq), CTCF occupancy or SMC3 occupancy at co-localized regions (**Fig. 2e-g; Supplemental Fig. S3f, g**). As expected, occupancy of SSBP3, an obligate co-factor for LDB1-dependent looping^22^, was diminished upon acute LDB1 depletion (**Fig. 2f, g**). In contrast, LMO2 was retained, consistent with a model in which LMO2 serves as a bridge between DNA-bound TFs (which by inference are unaffected by LDB1 loss). Hence, any observed transcriptional and architectural changes induced by LDB1 loss are likely directly attributable to a reduction in LDB1/SSBP3 looping activity instead of secondary effects from disrupted enhancer activity or the local chromatin environment.

### LDB1-dependent enhancer connectivity drives subtype-specific oncogenic transcription

To elucidate how distinct LDB1 chromatin occupancy profiles impact gene expression in different T-ALL subtypes, we performed transient transcriptome sequencing^54,55^ (TT-seq) upon acute LDB1 depletion in LOUCY^DH^ and KOPT K1^DH^ cells (**Fig. 3a; Supplemental Fig. S4a-e; Supplemental Tables 3-6**). Using log_2_(fold change [FC]) and *P_adj_* cutoffs of >|0.5| and <0.05, respectively, we detected 231 and 61 downregulated genes and 97 and 73 upregulated genes in LOUCY^DH^ and KOPT K1^DH^ cells, respectively, with minimal overlap between the two cell lines. LDB1-dependent genes, defined as those downregulated upon acute LDB1 depletion, in LOUCY^DH^ cells were notable for hematopoietic TFs and developmental oncogenes (e.g. *MYCN, ERG, MYB, HHEX, GATA2, LYL1*). Conversely, LDB1-dependent genes in KOPT K1^DH^ cells were associated with signal transduction and lymphocyte homeostasis (e.g. *DUSP6, BACH2, CD69, CD70, STAT4*). Nevertheless, pathway analysis suggested that these unique transcriptional circuits may still converge on common oncogenic pathways, such as MYC-driven transcription, KRAS signaling and JAK/STAT signaling (**Supplemental Fig. S4f**), perhaps explaining why LDB1 loss ultimately inhibits growth of T-ALL cells.

**Figure 3.**
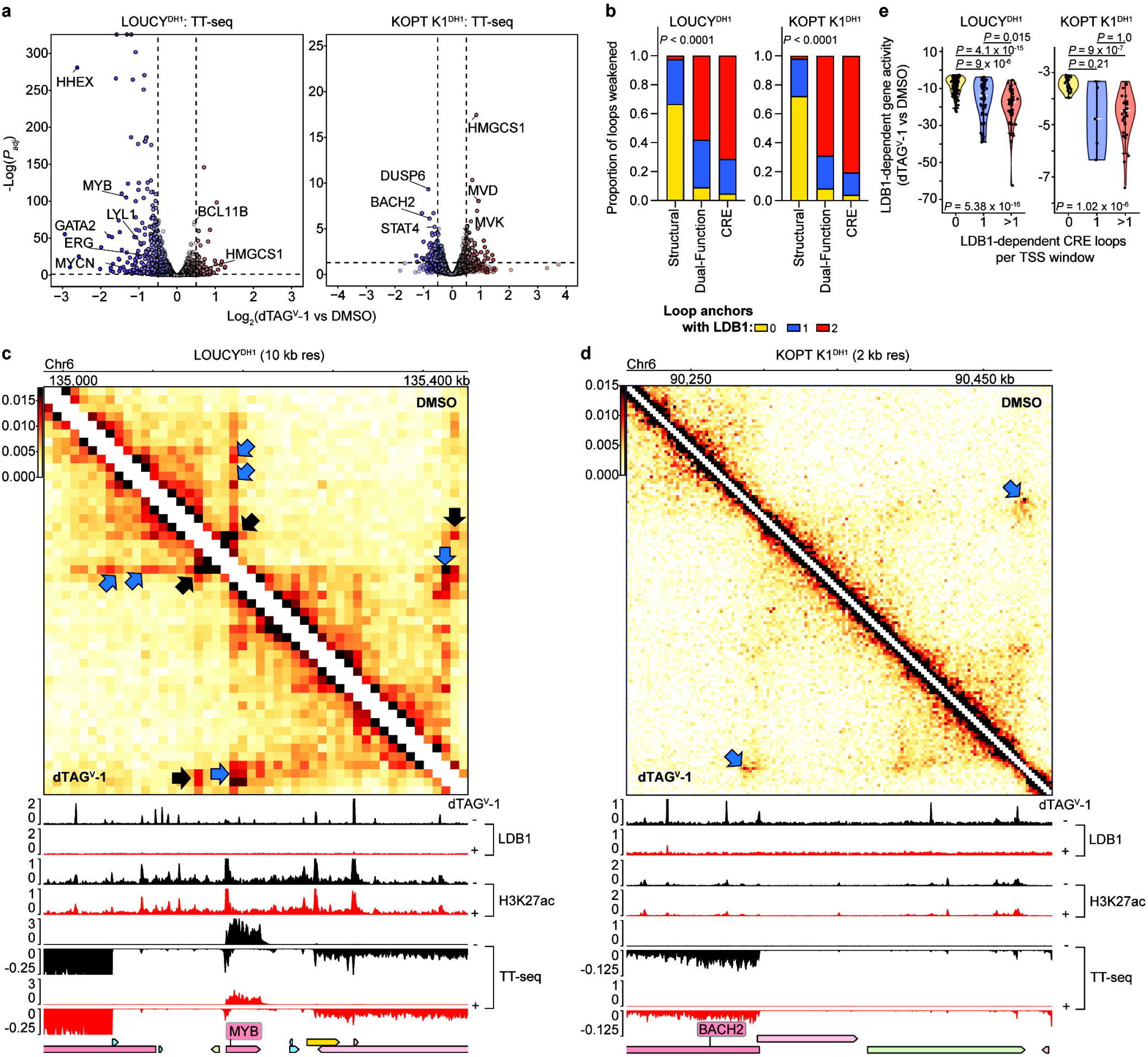
LDB1-dependent enhancer-promoter connectivity drives oncogenic transcriptional programs. **a,** Volcano plots of gene expression changes measured by TT-seq upon acute LDB1 depletion; relevant genes are denoted (blue, downregulated; red, upregulated). **b,** Weakened chromatin loops upon acute LDB1 depletion measured by Micro-C, stratified by loop class and number of LDB1-occupied anchors; *P*-values from Fisher’s exact test. **c,** 10 kb resolution Micro-C contact map of the *MYB* locus in LOUCY^DH^, depicting CRE loop changes upon 4-hour DMSO and dTAG^V^-1 treatments (blue arrow, weakened LDB1-dependent loops; black arrow, unchanged/strengthened LDB1-independent loops), with aligned LDB1 and H3K27ac from ChIP-seq and TT-seq tracks (black, DMSO; red, dTAG^V^-1). **d,** Similar to **c,** but depicting a 2 kb resolution contact map of the *BACH2* locus in KOPT K1^DH1^. **e,** Violin plots of LDB1-dependent gene expression changes (DESeq2 Wald statistic from TT-seq) upon acute LDB1 depletion, stratified by overlap with LDB1-dependent CRE loops; *P*-values from Kruskal-Wallis test with post hoc Dunn’s test. *N =* 4 biological replicates for TT-seq; *N = 3* pooled biological replicates for Micro-C per cell line and treatment. CRE, cis-regulatory element; TSS, transcription start site.

As the core function of LDB1 is to promote chromatin looping, we next investigated LDB1’s impact on chromatin architecture. We performed Micro-C^56–58^ after 4-hour LDB1 depletion and obtained approximately 1 billion valid contacts in each cell line and treatment (**Supplemental Table 7**). Acute LDB1 depletion had minimal impact to global chromatin compaction, compartmentalization or TAD boundary strength (**Supplemental Fig. S4g-j**), consistent with its predominant role in connecting CREs^21,22^. We used cooltools to map 38,752 and 30,724 chromatin loops in LOUCY^DH^ and KOPT K1^DH^, respectively, by merging loop calls across resolutions of 10, 5, and 2 kb to maximize sensitivity. We then quantified loop strength for each loop in each treatment using a log_2_(FC) cutoff of >|0.5| upon LDB1 depletion to define loops as weakened, strengthened or unchanged (**Supplemental Tables 8, 9**). Loops were then categorized as 1) structural loops (CTCF/SMC3 co-occupancy at both anchors with CRE at ≤1 anchor), 2) dual-function loops (CTCF/SMC3 co-occupancy and CREs at both anchors), 3) CRE loops (CREs at both anchors with CTCF/SMC3 co-occupancy ≤1 anchor) or 4) unclassified (not structural, dual-function or CRE). Using this schema, acute LDB1 depletion was more frequently associated with weakened CRE loops (18.0% of all CRE loops in LOUCY^DH^ and 22.1% of all CRE loops in KOPT K1^DH^) compared with dual-function or structural loops (*P* < 0.0001 for either cell line) (**Fig. 3b-d; Supplemental Fig. S5a, b**). As our Micro-C analysis was performed on pooled libraries, we confirmed that CRE loop changes were concordant between individual biological replicates (**Supplemental Fig. S5c**). Importantly, over 95% of weakened CRE loops had LDB1 present at one or both anchors. At higher resolutions, different loop calling methods can have substantial variability due to distinct prediction algorithms^59^. We therefore tested two additional loop prediction tools, mustache (merging 10, 5, and 2 kb resolution loop calls) and peakachu (merging 10 and 5 kb resolution loop calls) and quantified loop strength using cooltools. Although the absolute loop numbers differed, we similarly observed that a higher percentage of CRE loops were weakened upon LDB1 depletion than structural or dual-function loops (not shown). Our final analysis uses cooltools for consistency between loop calling and loop strength calculation.

We further stratified LDB1-dependent CRE loops, defined as weakened CRE loops with LDB1 present at ≥1 loop anchor, based on enhancer (E) and/or promoter (P) connectivity between loop anchors (**Supplemental Fig. S5d**). LDB1-dependent CRE loops were often E-E or mixed (E and P both found within the same loop anchor) contacts, suggesting LDB1 nucleates multi-enhancer hubs as well as co-regulated gene networks. Indeed, we frequently observed multiple LDB1-dependent CRE loops converging onto a single promoter such as at the *MYB* locus (**Fig. 3c; Supplemental Fig. S5e, f**).

To investigate the relationship between LDB1’s impact on transcription and chromatin architecture, we intersected CRE loop anchors with a 5 kb window surrounding the TSS of all active genes identified in our TT-seq data. Genes overlapping with LDB1-dependent loops were more likely to be downregulated upon acute LDB1 depletion (**Fig. 3e**). We also found that higher valency of LDB1-dependent loops was associated with a more potent effect on transcription, suggesting that some genes may be driven by multi-way or redundant enhancer contacts formed by LDB1. Loop formation at the *MYB, HHEX* and *MYCN* loci in LOUCY^DH^ cells serve as an illustrative example of this (**Fig. 3c; Supplemental Fig. S5e**): contacts between multiple LDB1-occupied CREs and the gene promoters were weakened upon LDB1 depletion (blue arrows), while nearby non-LDB1-dependent contacts were strengthened or unchanged (black arrows), ultimately resulting in potent downregulation of *MYB, HHEX* and *MYCN* gene expression. Examination of TT-seq tracks demonstrated that enhancer RNA signal at LDB1-occupied sites was largely preserved after LDB1 loss (**Fig. 3c, d**), orthogonally validating our observations with H3K27ac signal (**Fig. 2f, g; Supplemental Fig. S3f, g**) and supporting the concept that LDB1 serves as a looping factor with little impact on enhancer activity at least upon acute perturbation.

Together, our analysis of chromatin structural changes and nascent transcription upon acute LDB1 depletion indicates that LDB1-dependent spatial enhancer connectivity is required for the expression of critical oncogenes across T-ALL subtypes. Two technical limitations are worth noting. First, Micro-C likely underestimates LDB1-dependent contacts, particularly smaller loops and microcompartments that require ultra-high-resolution techniques to map, based on our previous work using Region Capture Micro-C^21^. For example, a contact between the known *GATA2* +9.5 intronic enhancer and its promoter can be visualized at 1 kb resolution (**Supplemental Fig. S5e**) but was missed by cooltools. Second, since it averages across populations, Micro-C cannot resolve whether contacts linking multiple CREs occur simultaneously or individually between alleles. Using Tri-C, however, we have previously shown that multi-way LDB1-dependent loops can occur, supporting the notion of multi-enhancer hubs^21^.

### LDB1 loss liberates enhancers to form promiscuous chromatin contacts

Although acute LDB1 depletion weakened approximately 20% of all CRE loops, we also observed a similar proportion of strengthened dual-function and CRE loops with LDB1 present at one or both anchors (**Supplemental Fig. S5a, b**). Because enhancer activity remained intact at LDB1-vacated sites, we reasoned that freed enhancers may be capable of interacting with new CREs. To test this, we asked whether the anchors of weakened LDB1-dependent CRE loops (“liberated anchors”) overlap with the anchors of newly gained or strengthened contacts (**Supplemental Fig. S6a**). We defined the new partner locus of these contacts as “destination anchors.” Using this framework, we found that rewiring of LDB1-dependent CRE loops was indeed common (**Supplemental Fig. S6b**), with 66.9% and 51.5% of LDB1-dependent CRE loops sharing an anchor with a strengthened loop in LOUCY^DH^ and KOPT K1^DH^, respectively. Although rewiring of non-LDB1 dependent loops was also observed, this may be attributable to false-negative LDB1 ChIP-seq peaks, as we have previously observed^22^, or secondary changes proximal to LDB1-dependent loop changes. Nevertheless, rewiring of LDB1-dependent loops was more frequently observed than with those lacking LDB1 (*P* = 0.0123 and *P* = 0.0382 for LOUCY^DH^ and KOPT K1^DH^, respectively). Therefore, the rewiring events we observe likely reflect a specific property of LDB1-dependent CRE loops, which maintain active chromatin marks upon LDB1 depletion and are therefore architecturally poised to establish new contacts.

Most de novo contacts originating from liberated anchors were directed toward other CREs (**Supplemental Fig. S6c**). To evaluate the transcriptional consequences of this rewiring, we intersected destination anchors with 5 kb windows surrounding TSSs of active genes in our TT-seq data and stratified genes by their expression change upon acute LDB1 depletion (**Supplemental Fig. S6d**). Rewiring of LDB1-dependent CRE loops was predominantly associated with transcriptional activation, accounting for 10.5% of upregulated genes in LOUCY^DH^ (*P* = 0.0169) and 12.3% of upregulated genes in KOPT K1^DH^ (*P* = 0.0288). There was no consistent transcriptional effect associated with rewiring of dual-function loops across both cell lines, potentially suggesting that structural elements may either have an inhibitory or negligible effect on enhancer rewiring or may block enhancer-mediated transcriptional activation. Although most upregulated genes could not be directly attributed to a rewired CRE loop, this may reflect Micro-C resolution and its limited ability to detect smaller loops as previously discussed.

### LDB1-dependent transcriptional networks are required for T-ALL growth

We assessed the direct functional consequences of LDB1 loss by devising a competitive cell proliferation assay using a CRISPRa approach to determine whether restoring select LDB1-dependent genes could rescue cell growth upon LDB1 depletion (**Fig. 4a**). We stably expressed CRISPRa in LOUCY^DH^ and KOPT K1^DH^ cells. We then selected genes with potential growth functionality based on LDB1-dependent looping, magnitude of transcriptional downregulation and known oncogenic roles, such as *MYCN, MYB* and *MEF2C*. We transduced CRISPRa cells with gRNA constructs to achieve a mixed population of gRNA-positive and negative cells and serially cultured them in the presence of DMSO or dTAG^V^-1 to continuously suppress LDB1. We measured the proportion of gRNA-positive to gRNA-negative (untransduced) cells in relation to cells with a non-targeting control (NTC) gRNA. Cells harboring NTCs failed to expand upon LDB1 loss as expected. In LOUCY^DH^, even partial restoration of *MYCN, MYB* and *MEF2C* resulted in improved cell growth (**Fig. 4b, c)**. In KOPT K1^DH^, restoration of *DUSP6* and *STAT4* provided only a modest growth advantage which did not reach statistical significance (**Fig. 4d, e**). Nonetheless, no single gene was able to completely rescue cell growth, indicating that LDB1 dependency is encoded across a broader transcriptional network rather than a single oncogenic node.

**Figure 4.**
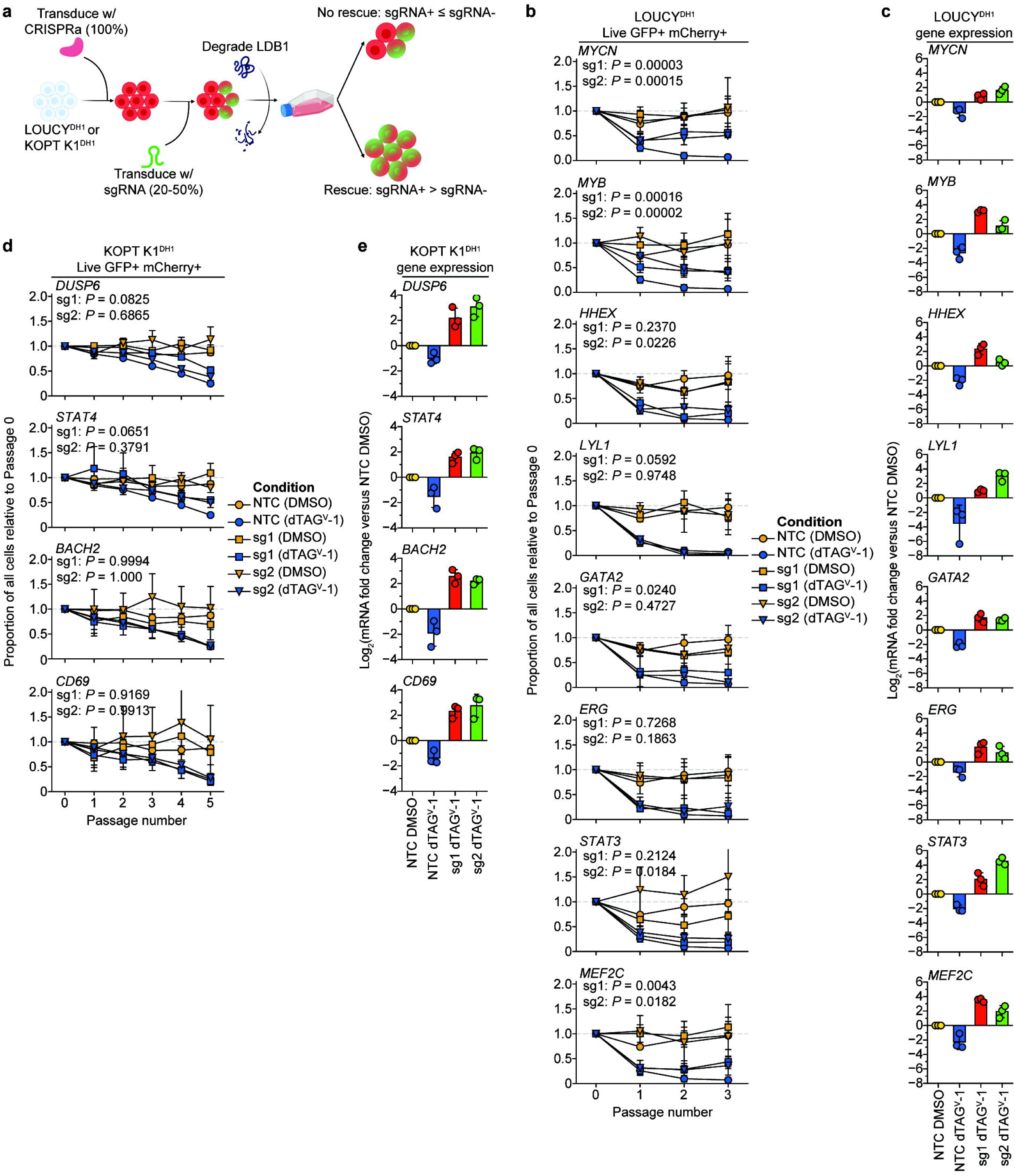
LDB1-dependent genes are required for T-ALL survival. **a,** Schematic of CRISPRa (dCas9-VPR) competitive cell proliferation assay. **b,** Proportion of live GFP+ mCherry+ cells shown for non-targeting control (NTC), gene promoter-targeting CRISPRa sgRNAs in LOUCY^DH1^; values normalized to Passage 0 for each sgRNA; *P*-values from Dunnett’s test (two-sided, FWER < 0.05) comparing each group to NTC using pooled log-transformed replicate values from all non-Passage 0 passages. **c,** mRNA expression for CRISPRa gene targets shown from RT-qPCR in LOUCY^DH1^ with or without LDB1 depletion; log_2_-transformed values for dTAG^V^-1 treatments are shown relative to NTC/DMSO control. **d,** similar to **b,** but depicting KOPT K1^DH1^ CRISPRa cell competition assay. **e,** similar to **c,** but depicting KOPT K1^DH1^ CRISPRa RT-qPCR. *N =* 3 biological replicates per cell line, sgRNA and treatment.

### LDB1-dependent chromatin loops are conserved in primary T-ALL in a subtype-specific manner

To evaluate whether LDB1-dependent loops identified in our cell line models represent those found in primary disease, we compared our Micro-C findings with publicly available H3K27ac HiChIP data from patients with T-ALL (*N =* 29) and healthy hematopoietic controls (CD34+ HSPCs and double-positive [DP] thymocytes, *N =* 1 each)^36,47^. We called loops in primary T-ALL HiChIP datasets using FitHiChIP and filtered them based on the presence of H3K27ac peaks at both loop anchors, fitting our definition of dual-function or CRE loops. After removing samples with poor H3K27ac capture (leaving *N =*17) and removing duplicative overlaps (e.g. one Micro-C loop overlapping with two HiChIP loops), a high degree of LDB1-dependent dual-function and CRE loops from both LOUCY^DH^ (42.7% ± 14.8%) and KOPT K1^DH^ (52.1% ± 16.7%) Micro-C were detectable in primary T-ALL HiChIP datasets (**Fig. 5a, b; Supplemental Table 10**), with most variance driven by 3 samples of sub-optimal quality (<15,000 HiChIP loops called). This remarkably high overlap is likely still an underestimate given the different techniques used, different loop calling algorithms and the relative sparsity of some of the primary cell data. To assess subtype specificity, we first compared the ratio of LDB1-dependent CRE loop overlap between ETP/near-ETP and non-ETP HiChIP samples for each cell line (**Fig. 5c**). LDB1-dependent CRE loops from LOUCY^DH^ showed preferential overlap with ETP/near-ETP samples relative to non-ETP samples, while the converse was observed for KOPT K1^DH^, mirroring the known differentiation states of each cell line.

**Figure 5.**
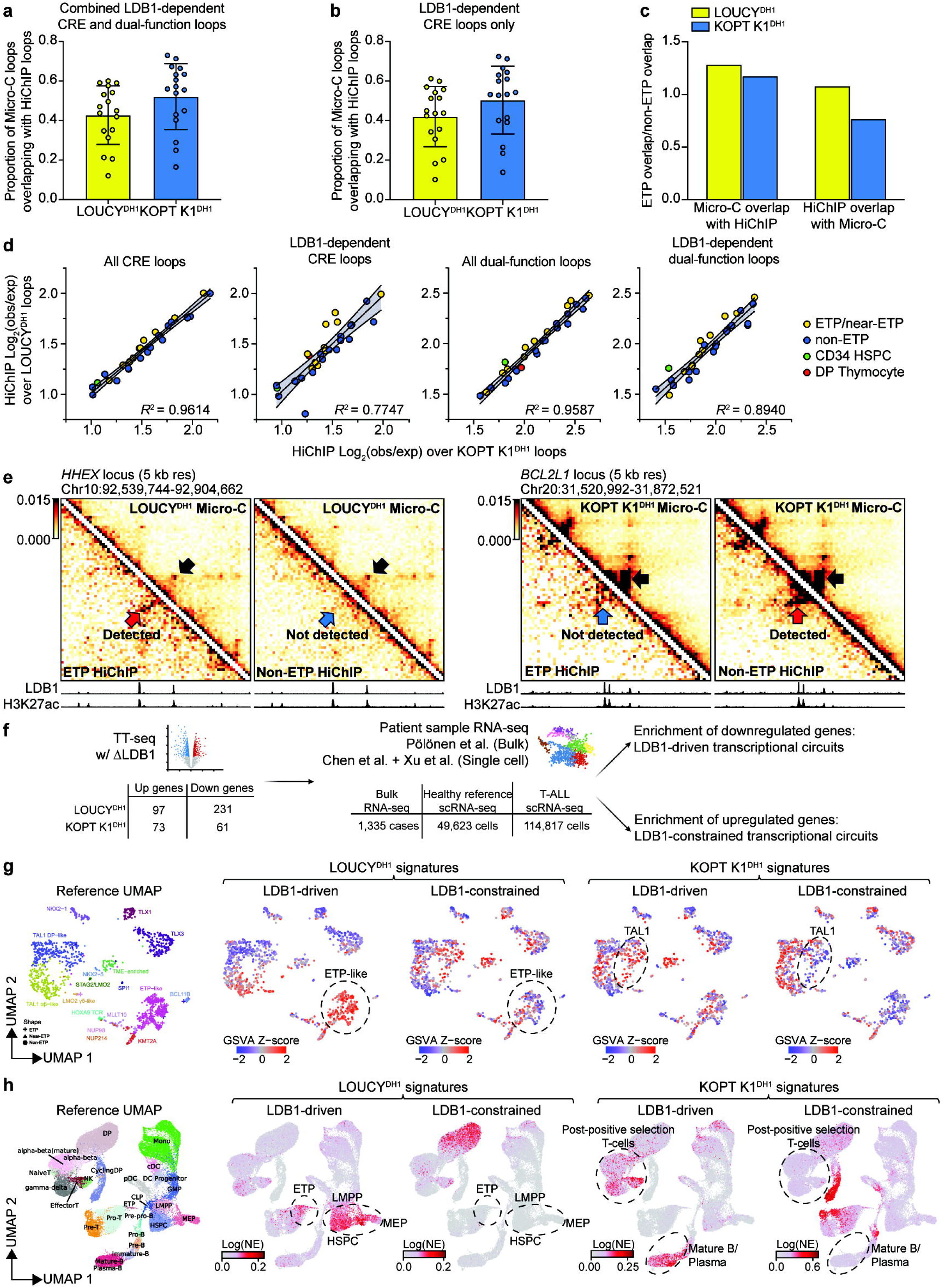
LDB1-dependent chromatin looping and transcription defines subtype-specific T-ALL transcriptional states. **a,** Proportion of pooled LDB1-dependent CRE/dual-function loops called by cooltools from LOUCY^DH1^ and KOPT K1^DH1^ Micro-C overlapping with loops called by FitHiChIP from primary H3K27ac HiChIP datasets (*N* = 17), filtered on HiChIP loops with H3K27ac present at both anchors and duplicative overlaps removed. **b,** Similar to **a**, but filtered on CRE loop calls only from Micro-C. **c,** Ratio of overlap frequencies between ETP and non-ETP H3K27ac HiChIP samples based on proportion of Micro-C loops detected in H3K27ac HiChIP datasets and vice versa. **d,** Scatter plots depicting loop strength for H3K27ac HiChIP samples (*N* = 29) over CRE or dual-function loops defined by LOUCY^DH1^ versus KOPT K1^DH1^ Micro-C; best-fit linear regression (line) with 95% CI (shaded) and *R*^2^ shown. **e,** 5 kb resolution contact maps comparing detection of chromatin loops from LOUCY^DH1^ and KOPT K1^DH1^ Micro-C to primary T-ALL H3K27ac HiChIP at the *HHEX* and *BCL2L1* loci, respectively (black arrow, LDB1-dependent loop in Micro-C; blue arrow, absent/undetected loop in HiChIP; red arrow, detected loop in HiChIP), with aligned LDB1 and H3K27ac ChIP-seq tracks from DMSO-treated cells. **f,** Schematic of TT-seq comparison to primary T-ALL transcriptomic datasets, including sample sizes. **g,** UMAP plots depict a primary T-ALL whole transcriptome sequencing reference dataset^36^, with ETP status, molecular subtype annotations and projections of GSVA Z-scores from LOUCY^DH1^ or KOPT K1^DH1^ TT-seq gene sets; each point represents one patient (*N =* 1,335); subtypes with LDB1-driven transcriptional circuits are outlined (see **Supplemental Fig. S7a-d** for GSVA summary data). **h,** UMAP plots depict a healthy pediatric hematopoiesis scRNA-seq reference dataset^60,61^, with hematopoietic cell type annotations and projections of normalized expression values from LOUCY^DH1^ or KOPT K1^DH1^ TT-seq gene sets; each point represents one cell (*N =* 49,623); cells with LDB1-driven transcriptional circuits are outlined. CRE, cis-regulatory element; ETP, early thymic precursor; HSPC, hematopoietic stem and progenitor cell; DP, double-positive thymocyte.

Several of the primary HiChIP samples had adequate sequencing depth despite sub-optimal H3K27ac capture, allowing them to be used for an orthogonal loop strength analysis unbiased by H3K27ac presence at loop anchors. Therefore, we calculated observed/expected loop strength for 27 T-ALL (2 excluded due to low quality) and 2 control HiChIP datasets over Micro-C-defined CRE or dual-function loop coordinates (**Fig. 5d**), which confirmed this subtype-specific pattern through an independent metric: all CRE loops from either cell line showed near-identical strength across all patient HiChIP samples regardless of ETP status (*R²* > 0.9), while LDB1-dependent CRE loops showed preferential alignment with ETP/near-ETP HiChIP samples for LOUCY^DH^ and non-ETP samples for KOPT K1^DH^. This subtype-specific skewing was more pronounced in T-ALL samples than in normal hematopoietic controls, suggesting that a subset of LDB1-dependent CRE loops may be formed de novo or selectively strengthened in T-ALL rather than simply reflecting developmental arrest. As an illustrative example, at the *HHEX* locus, the LDB1-dependent loop is stronger in ETP HiChIP data compared with non-ETP HiChIP data (**Fig. 5e, left**, compare red and blue arrows). Conversely, at the *BCL2L1* locus, the LDB1-dependent loop is stronger in non-ETP HiChIP data compared with ETP HiChIP data (**Fig. 5e**, **right**, compare blue and red arrows).

In sum, we observed a remarkable similarity between CRE loops in cell lines and those in primary T-ALL samples, validating the former as model system for mechanistic studies that are impossible in the latter.

### LDB1-dependent transcriptional circuitry defines T-ALL identities

To examine whether LDB1-dependent regulatory connectivity is reflected in transcriptional output between primary T-ALL subtypes, we leveraged recently published transcriptomic data^36,60,61^ from patients with T-ALL enrolled in AALL0434 (<u>NCT00408005</u>), an international phase 3 randomized Children’s Oncology Group (COG) trial, as well as normal hematopoietic controls (**Fig. 5f**). We operationally defined genes downregulated upon acute LDB1 depletion as “LDB1-driven” circuits and genes upregulated upon LDB1 depletion as “LDB1-constrained”. Projecting these gene sets onto a whole transcriptome sequencing atlas from 1,335 patients clustered by molecular subtype (**Fig. 5g**) revealed that LDB1-driven circuits from LOUCY^DH^ were preferentially enriched in ETP-like ALL and NUP214-rearranged T-ALL, implicating LDB1-dependent chromatin architecture as a key enforcer of the ETP-like transcriptional program, a high-risk identity that spans multiple molecular subtype boundaries^36^ and lacks a clear mechanistic basis (**Supplemental Fig. S7a, b**). Conversely, LDB1-driven circuits from KOPT K1^DH^ were enriched in a subset of TAL1 DP-like T-ALL, a non-ETP ALL characterized by chromosomal rearrangements involving the TAL1 transcription factor^36^ (**Supplemental Fig. S7c, d**). However, a subset of patients with the TAL1 DP-like subtype lacks *TAL1* expression and instead harbors *LMO2* rearrangements or enhancer hijacking events^36^, indeed emulating the transcriptional and genetic state of KOPT K1 cells (**Supplemental Table 1**). In both cases, LDB1-constrained circuits corresponded to transcriptional programs of alternative molecular subtypes, indicating that LDB1 actively enforces subtype-specific transcriptional identity while restricting plasticity toward other leukemic states.

Comparison of LDB1-driven and LDB1-constrained gene sets against single-cell RNA-seq (scRNA-seq) reference atlases from healthy pediatric hematopoiesis (*N =* 49,623 cells) and primary T-ALL (*N =* 114,817 cells) recapitulated these findings at single-cell resolution (**Fig. 5h**). LDB1-driven genes from LOUCY^DH^ were enriched in bone marrow progenitor-like (BMP-like^60,61^) cell states (**Supplemental Fig. S7e-g)** while those from KOPT K1^DH^ aligned with more mature T- and B-cell states (**Supplemental Fig. S7e, f**). LDB1 loss in each context was associated with a reciprocal shift toward the opposing maturational pole, consistent with LDB1 actively constraining differentiation state in both directions and in a manner conserved across healthy hematopoietic and primary T-ALL reference datasets.

### LDB1-driven transcriptional signatures are associated with high-risk subgroups within TAL1-related T-ALL

Molecular identities of T-ALL subtypes are closely linked with clinical outcomes^36^, and our findings indicate that LDB1 is critical for enforcing such identities. Therefore, we examined whether enrichment of LDB1-driven signatures served as a prognostic indicator. Enrichment of the LOUCY^DH^ LDB1-driven signature was associated with significantly worse survival across all patients with T-ALL (*N =* 1,335; **Fig. 6a**). Within the ETP-like subgroup (*N =* 240), this signature lost independent prognostic significance, which is mechanistically predicted by our model: if LDB1-driven programs define the ETP-like transcriptional state broadly, the signature should distinguish ETP-like from other T-ALL subtypes rather than stratify risk within a group where high-risk features are already uniform. Multivariable analysis supports this interpretation, as high-risk features associated with ETP-like ALL, such as measurable residual disease, decrease the prognostic value of this signature (**Fig. 6b; Supplemental Table 11**). The KOPT K1^DH^ LDB1-driven signature did not stratify outcomes across all T-ALL patients or ETP-like ALL patients (**Fig. 6c, d; Supplemental Table 11**).

**Figure 6.**
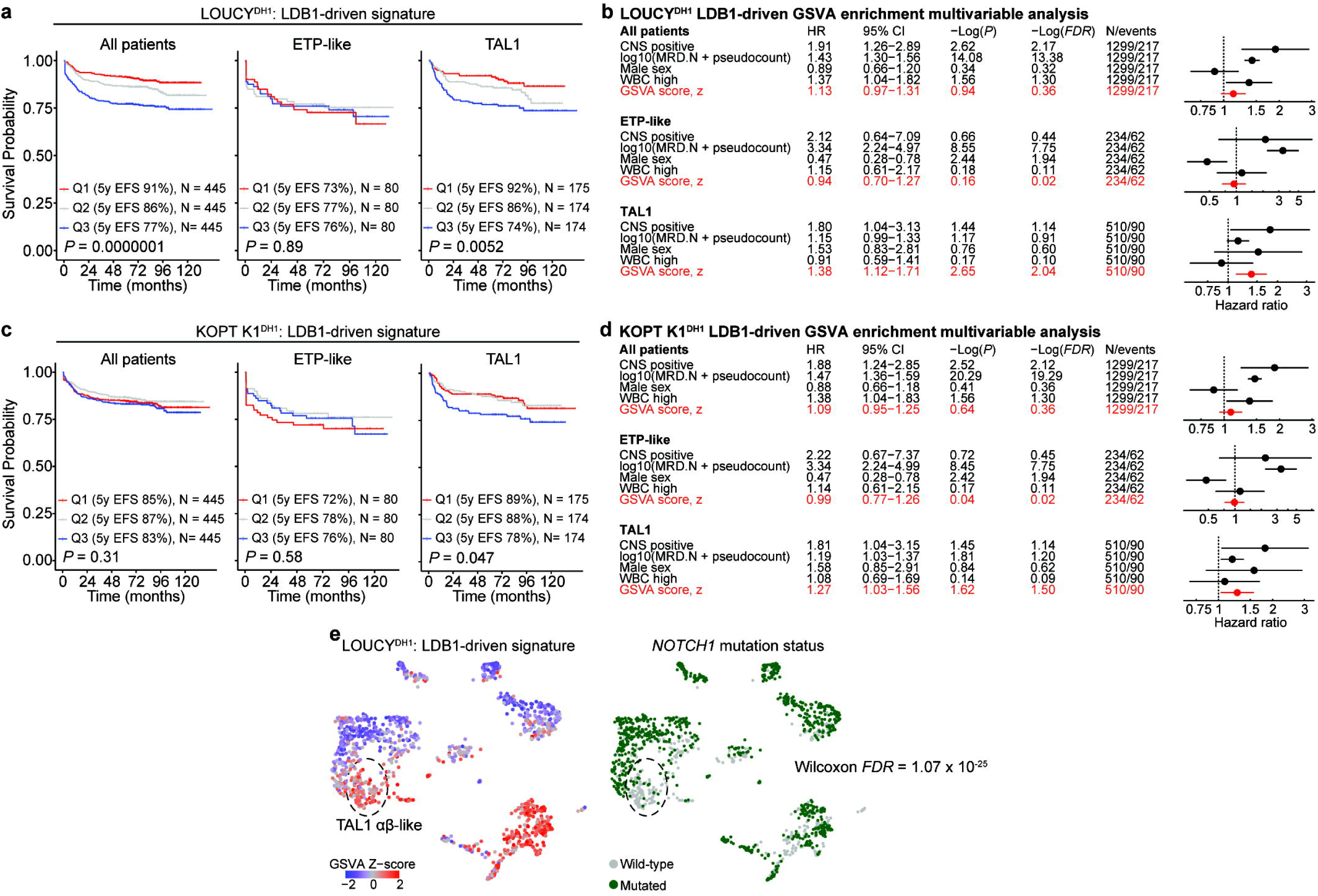
LDB1-dependent transcriptional profiles are associated with high-risk subtypes of T-ALL. **a,** Kaplan-Meier event-free survival (EFS) curves for all patients with T-ALL (*N =* 1,335), ETP-like ALL (*N =* 240) and TAL1-related T-ALL (*N* = 523), stratified by LOUCY^DH1^ LDB1-driven gene signature enrichment (Q1/red, lowest tertile; Q2/grey, middle tertile; Q3/blue, highest tertile); *P*-values from log-rank test. **b,** Forest plots depicting multivariable analyses for LOUCY^DH1^ LDB1-driven GSVA enrichment with clinical covariates based on complete case analysis of all patients (*N* = 1,299), ETP-like ALL (*N* = 234), and TAL1-related T-ALL (*N* =510); hazard ratio (HR) >1 indicates higher risk of EFS events; *N* for each comparison group denoted in panel based on complete case analysis. **c,** Similar to **a,** but depicting Kaplan-Meier EFS curves stratified by KOPT K1^DH1^ LDB1-driven gene signature enrichment. **d**, Similar to **b**, but depicting KOPT K1^DH1^ LDB1-driven GSVA enrichment multivariable analysis. **e,** UMAP plots depict projection of the LOUCY^DH1^ LDB1-driven GSVA Z-scores and *NOTCH1* mutation status onto primary T-ALL whole transcriptome sequencing datasets (see Fig. 5g for reference atlas); each point represents one patient (*N =* 1,335); TAL1 αβ-like samples with *NOTCH1* wild-type status are outlined (see **Supplemental Fig. 7a, b** for GSVA summary data); association of GSVA enrichment with *NOTCH1* wild-type status calculated by two-sided Wilcoxon rank-sum test (see **Supplemental Table 12**). GSVA, gene set variation analysis; CNS, central nervous system; MRD, measurable residual disease; WBC, white blood cell count; ETP, early thymic precursor; EFS, event-free survival; CI, confidence interval.

Notably, we observed that both LOUCY^DH^ and KOPT K1^DH^ LDB1-driven signatures were enriched in distinct subsets of TAL1-related leukemias (TAL1 αβ-like and TAL1 DP-like, respectively) and were predictive of worse outcomes after correction for other clinical covariates (**Fig. 6b, d; Supplemental Fig. S7a-d; Supplemental Table 11**). Patients with LOUCY^DH^ LDB1-driven signature enrichment were markedly depleted for *NOTCH1* mutations (*FDR* = 1.07 X 10 from two-sided Wilcoxon rank-sum test), particularly those in the TAL1-related subgroup (**Fig. 6e, Supplemental Table 12**). Together, these findings suggest that LDB1-dependent transcriptional circuit engagement contributes to disease aggressiveness within the heterogeneous TAL1-related T-ALL and may define a novel high-risk molecular subgroup characterized by *NOTCH1*-wild-type disease.

### Enhancer rewiring drives mevalonate pathway addiction and new metabolic dependencies

To determine whether enhancer rewiring events create any exploitable dependencies, we examined LDB1-constrained genes from our TT-seq experiments and found several cholesterol biosynthesis/mevalonate pathway genes (*HMGCS1, MVK, MVD*) upregulated in both LOUCY^DH^ and KOPT K1^DH^ upon LDB1 depletion (**Fig. 3a**; **Fig. 7a**). Analysis of Micro-C contact maps at these loci revealed that some LDB1-anchored CRE loops flanking the *HMGCS1* and *MVD* promoters were weakened upon LDB1 depletion (**Fig. 7b-e; Supplemental Fig. S8a, b**). Consistent with our proposed rewiring mechanism, enhancer activity at the vacated loop anchors remained intact, and several of the elements involved in these loops instead formed strengthened contacts with the *HMGCS1* and *MVD* promoters.

**Figure 7.**
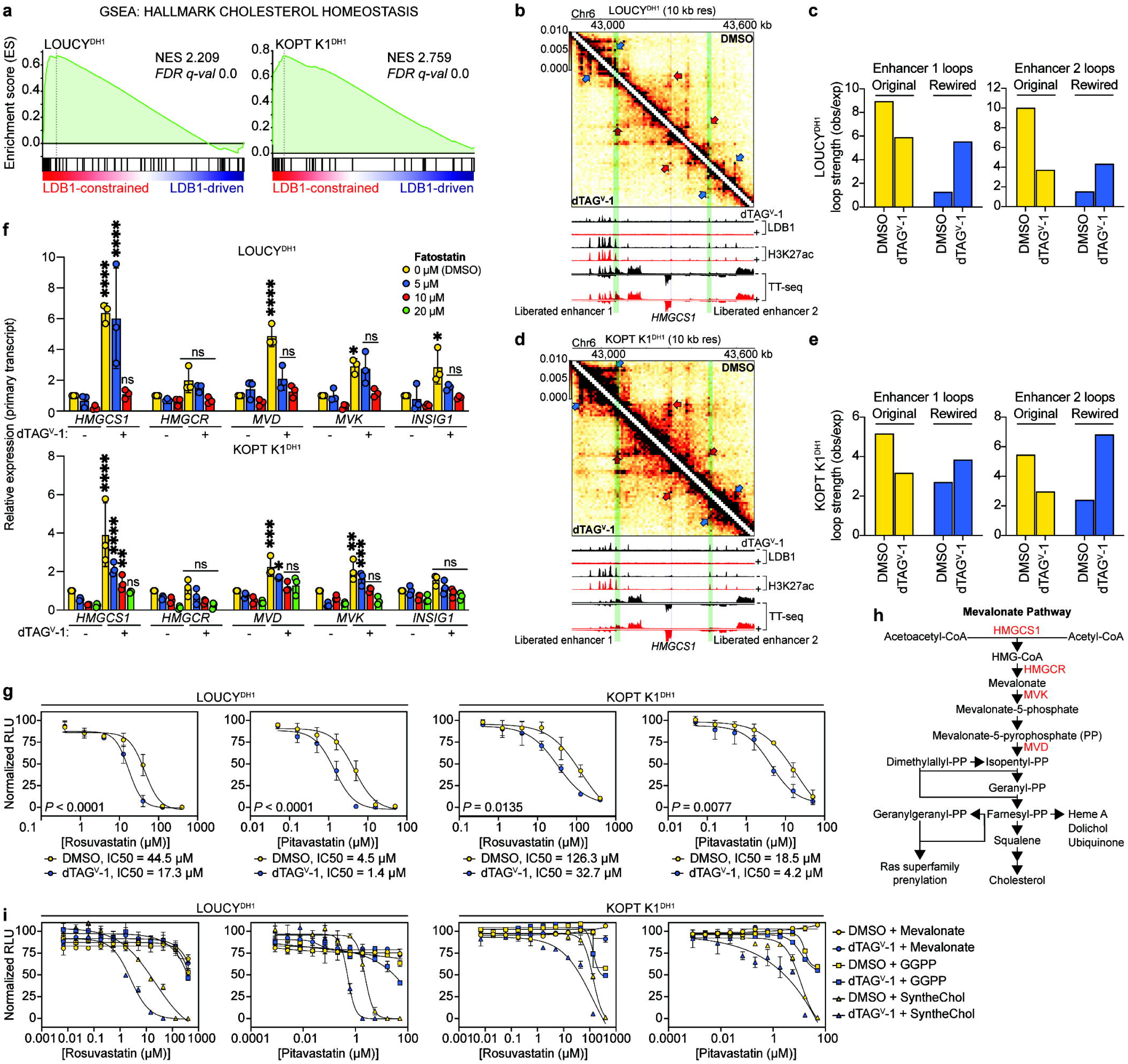
Chromatin loop rewiring upon LDB1 depletion drives mevalonate pathway dependency in T-ALL. **a,** GSEA for MSigDB Hallmark Cholesterol Homeostasis gene set in TT-seq data from LOUCY^DH1^ and KOPT K1^DH1^ upon acute LDB1 depletion. **b,** 10 kb resolution Micro-C contact map of the *HMGCS1* locus in LOUCY^DH1^, depicting CRE loop rewiring events upon 4-hour DMSO and dTAG^V^-1 treatments (blue arrow, weakened loops; red arrow, strengthened loops; purple shading, *HMGCS1* promoter; green shading, liberated anchors/enhancers), with aligned LDB1 and H3K27ac ChIP-seq and TT-seq tracks (black, DMSO; red, dTAG^V^-1). **c,** Observed/expected loop strength for rewired loops in LOUCY^DH1^ corresponding to panel **b** (for a shared enhancer/anchor: yellow, weakened/original loop; blue: strengthened/rewired loop). **d,** Similar to **b,** but depicting KOPT K1^DH1^. **e,** Similar to **c,** but depicting KOPT K1^DH1^ loop strengths corresponding to panel **d**. **f,** Relative primary transcript expression of mevalonate pathway genes shown from RT-qPCR in LOUCY^DH1^ and KOPT K1^DH1^ upon acute LDB1 depletion with or without fatostatin pre-treatment; expression levels normalized to DMSO/DMSO; *P*-values from two-way ANOVA with post hoc Dunnett’s test comparing dTAG^V^-1/fatostatin combinations to DMSO/DMSO per gene. **g,** Dose-response curves for combinatorial LDB1 depletion with statin treatment in LOUCY^DH1^ and KOPT K1^DH1^; viability quantified as CellTiter-Glo 2.0 relative luminescence units (RLU) and normalized to statin-negative controls per DMSO or dTAG^V^-1 condition; *P*-values from extra sum-of-squares F test. **h,** Summary of mevalonate pathway with metabolites (black text) and relevant genes/enzymes (red text). **i,** Similar to **g**, dose-response curves depict combinatorial LDB1 depletion, statin treatment, mevalonate pathway metabolite supplementation. *N = 3* biological replicates for all experiments. *NES*, normalized enrichment score; FDR, false discovery rate; IC50, half-maximal inhibitory concentration; GGPP, geranylgeranyl pyrophosphate.

Sterol-responsive binding element proteins (SREBPs) are master TFs for mevalonate pathway gene expression and are activated through proteolytic cleavage^62^. Fatostatin tethers SREBPs to the endoplasmic reticulum, inhibiting their transactivation^63^. Indeed, fatostatin pre-treatment of LOUCY^DH^ and KOPT K1^DH^ cells attenuated SREBP target gene expression upon acute LDB1 depletion in a dose-dependent manner (**Fig. 7f**). This suggests that SREBP-driven transcription is required for basal mevalonate pathway expression, but that neo-contacts from rewired enhancers substantially augment transcriptional output upon LDB1 depletion.

To verify that these findings are not artifacts of the dTAG^V^-1 system, we treated parental LOUCY and KOPT K1 cells with dTAG^V^-1 and did not observe similar transcriptional changes (**Supplemental Fig. S9a**). AKT/mTOR signaling has also been implicated in altered mevalonate pathway regulation in T-ALL^64^, but we did not observe significant changes to AKT, mTOR or S6 phosphorylation upon 4-hour LDB1 depletion (**Supplemental Fig. S9b**), suggesting that mevalonate pathway upregulation is a bona fide transcriptional consequence of LDB1 loss rather than a secondary effect of acute signaling dysregulation.

To test the functional consequences of LDB1-dependent activation of SREBP targets, we treated LOUCY^DH^ and KOPT K1^DH^ cells with the HMGCR inhibitors rosuvastatin or pitavastatin with or without concomitant dTAG^V^-1 (**Fig. 7g**). LDB1 depletion led to a 2-5-fold sensitization to statin therapy for both T-ALL cell lines, which we orthogonally validated via CRISPR/Cas9-mediated LDB1 knockout (**Supplemental Fig. S9c**). LDB1 knockout potentiated statin sensitivity in two additional LDB1-dependent cell lines (JURKAT and PF-382) but not in LMO1/2-deficient SUP-T1 cells, supporting that this vulnerability is specific to LDB1/LMO-dependent activity (**Supplemental Fig. S9c**).

Sustained LDB1 loss via CRISPR/Cas9 resulted in broader induction of mevalonate pathway genes, including *HMGCR* and *SREBP2* itself (**Supplemental Fig. S9d**), suggesting that prolonged LDB1 depletion drives an addiction to this pathway that can be therapeutically exploited with statins. To examine whether this relationship between LDB1 and mevalonate pathway suppression is conserved, we analyzed RNA-seq^31^ from murine double-negative 4 (DN4) thymocytes and found that *Lmo2* overexpression increased expression of putative LDB1/LMO2 target oncogenes, while simultaneously suppressing expression of mevalonate pathway genes (**Supplemental Fig. S9e**). *Rag1*-*Cre*-inducible *Ldb1* deletion in this model transcriptionally de-repressed *Hmgcs1*, *Mvd*, *Hmgcr* and *Srebp2* but not *Srebp1*, recapitulating our observations in human T-ALL cell lines and indicating that mevalonate pathway upregulation induced by LDB1 loss in T-ALL is conserved across species.

The mevalonate pathway produces multiple byproducts critical for cellular homeostasis (**Fig. 7h**). Previous work^64^ demonstrated that intracellular cholesterol pools regulate oncogenic signaling and drive a selective mevalonate pathway dependency in ETP ALL over non-ETP ALL. However, we observed that LDB1 loss sensitizes non-ETP ALL cell lines to statins as well. To determine whether this reflects a metabolic switch distinct from the cholesterol biosynthesis branch of the mevalonate pathway, we performed metabolite supplementation experiments with multiple mevalonate pathway byproducts (**Fig. 7i**). Cholesterol supplementation did not rescue statin sensitization, whereas mevalonate, an upstream metabolite of this pathway, or geranylgeranyl pyrophosphate (GGPP), an alternative branch of the mevalonate pathway, completely rescued statin-induced cell death and sensitization by LDB1 depletion.

Together these findings indicate that LDB1 loss elicits a targetable dependency on the GGPP branch of the mevalonate pathway rather than cholesterol biosynthesis per se.

## Discussion

Most prior studies specifically investigating the relationship between genome folding and cancer have been largely descriptive or have focused on the CTCF/cohesin machinery^36,45–47,65–75^. Since a substantial fraction of enhancer-promoter loops is independent of CTCF/cohesin^21,22,76–83^, this leaves unexplored the role of other essential mediators of spatial enhancer-enhancer and enhancer-promoter connectivity. Here, by acutely depleting the architectural factor LDB1 in T-ALL, we demonstrate that a single looping factor can simultaneously enforce subtype-specific oncogenic transcriptional programs and constrain alternative cell states. LDB1 loss liberates enhancers to rewire their connectivity to create new therapeutic vulnerabilities, establishing chromatin loop architecture as a bidirectional enforcer of leukemic identity. More broadly, our study demonstrates that perturbation of a chromatin architectural factor can expose dependencies that are otherwise hidden under homeostasis. Specifically in T-ALL, mutations creating new TF binding sites and structural variants activating subtype-defining TFs are as prevalent as coding driver mutations^36^. We propose a model in which these noncoding alterations collectively specify leukemic transcriptional programs by remodeling the TF landscape in a subtype-specific manner, while LDB1-dependent enhancer connectivity serves as the architectural enforcer of these programs.

A principal discovery of our study is that LDB1 not only maintains chromatin contacts among enhancers and oncogenes, but that it simultaneously constrains enhancers from activating gene expression programs which otherwise activate novel dependencies. This model is supported by our observation that 1) LDB1 loss has minimal impact on enhancer activity itself, that 2) unconstrained enhancers retain the potential to physically pair with illegitimate target genes and activate their expression, such as those involving mevalonate pathway genes, and 3) that interference with such newly activated genes can lead to cancer cell growth defects. The molecular mechanisms that spatially connect LDB1-vacated enhancers with new partners remain unknown but will be important to elucidate in future studies as it may enable predictions of new vulnerabilities across disease contexts with implications for understanding and potentially targeting T-ALL and related malignancies.

Our study also identified a molecularly distinct and clinically aggressive subset of TAL1-related T-ALL that is not captured by current risk stratification. Specifically, a subgroup of patients with TAL1-related T-ALL who lack canonical *NOTCH1* mutations harbor high enrichment of the LOUCY^DH^ LDB1-driven transcriptional signature and have significantly worse clinical outcomes. *NOTCH1* mutations are among the most prevalent and prognostically favorable alterations in T-ALL^36,43,44,84^, and their mutual exclusivity with LDB1 signature enrichment in this subgroup suggests that LDB1-dependent transcriptional circuitry drives leukemogenesis in the absence of canonical *NOTCH1*-driven oncogenic programs. Most of these patients harbor *LMO1/2* overexpression^36^, though the specific TFs that recruit LDB1/LMO2, thereby driving the transcriptional features of this subgroup, remain to be determined. Our data provide a mechanistic rationale for prioritizing LDB1-dependent programs as a defining feature and potential therapeutic target in *NOTCH1*-wild-type TAL1-related disease.

An unexpected finding was alignment of KOPT K1^DH^ LDB1-driven programs with B-cell transcriptional states, suggesting that LDB1-dependent looping may contribute to lymphoid oncogenesis beyond limiting T-cell maturation. LMO2 and its paralogs are dysregulated across multiple cancer types^85^, including diffuse large B-cell lymphoma^86–88^, where any architectural role of LDB1 remains largely unknown. Whether subtype-specific LDB1 occupancy patterns enforce transcriptional identities and create analogous survival dependencies in B-cell or other LMO-dysregulated malignancies is an important open question.

Mevalonate pathway activation upon LDB1 loss provides a proof-of-concept for chromatin rewiring as a source of therapeutic vulnerability. The observation that GGPP, but not cholesterol, rescues statin sensitization implicates the protein prenylation branch of the mevalonate pathway as the relevant downstream effector. Prenylation of Ras- and Rho-family GTPases by geranylgeranyl transferases is required for their membrane anchoring and oncogenic signaling^89,90^, and disruption of this modification has been explored as a therapeutic strategy in other cancer contexts^91^. Farnesyltransferase and geranylgeranyl transferase inhibitors targeting related branches of this pathway are in clinical development for other malignancies, and our findings raise the possibility that such agents could be combined with strategies to disrupt the LDB1/LMO2 complex. More broadly, statin sensitization was consistent across ETP and non-ETP subtypes and validated in four independent cell lines, implicating a generalizable relationship between LDB1/LMO-dependent looping and mevalonate pathway homeostasis that is mechanistically distinct from the cholesterol-driven vulnerability previously described in ETP ALL^64^.

Collectively, our findings provide a paradigm for understanding cancer subtype heterogeneity through the simultaneous activation and repression of defining transcriptional pathways via spatial organization of chromatin. While our model focuses on active enhancer connectivity as the primary driver of leukemic identity, repressive chromatin elements may also contribute to LDB1-constrained transcriptional programs, and future studies incorporating these features will further refine our understanding of how 3D genome architecture enforces cancer cell identity. As chromatin architectural regulators become better characterized, targeting loop-dependent vulnerabilities like those described here may offer new therapeutic avenues for cancers driven by dysregulated enhancer connectivity.

## Methods

### Engineering LDB1-dTAG-HaloTag isogenic cell lines

LOUCY and KOPT K1 LDB1-dTAG-HaloTag (LOUCY^DH^ and KOPT K1^DH^) cell lines were engineered using CRISPR-mediated homology-dependent repair. A donor template was designed to insert dTAG-HaloTag in-frame with the 3’ end of LDB1 using a flexible (G)_4_S linker to replace the endogenous stop codon. The donor template had 900 bp arms flanking the dTAG-HaloTag insert, sharing 5’ and 3’ homology with LDB1; homology arms were amplified from LOUCY or KOPT K1 genomic DNA to account for any potential sequence variants between the cell lines. The donor template was cloned into a transfer plasmid using the In-Fusion Snap Assembly kit (Takara Bio) and linearized using Platinum SuperFi II DNA Polymerase (Invitrogen) and gel-purified using the QIAquick Gel Extraction Kit (Qiagen). We designed two Alt-R sgRNAs (IDT) targeting PAM sites near the LDB1 stop codon (see **Supplemental Table 13).** Alt-R sgRNAs were complexed with Alt-R S.p. HiFi Cas9 V3 (IDT) at 220 pmol:220 pmol in a 10 µL reaction for 20 minutes at room temperature. Each ribonucleoprotein complex was mixed with 3.3 µg of the linearized dsDNA donor template, 4 µM Alt-R Electroporation Enhancer (IDT) and SE Cell Line Nucleofection Solution plus Supplement 1 (Lonza). LOUCY and KOPT K1 cells (1e6 each) were centrifuged at 125g for 10 minutes and resuspended in the nucleofection solution, transferred to a cuvette and nucleofected using a Lonza 4D-Nucleofector system with program CL-120. Cells were then immediately quenched with pre-warmed antibiotic-free media, transferred to a 12-well plate, supplemented with Alt-R HDR Enhancer V2 (IDT) and incubated for 16 hours prior to washing and resuspending into fresh media. After 24-48 hours, we confirmed HaloTag expression by adding 40 nM Janelia Fluor 646 HaloTag Ligand (Promega) directly in cell culture and incubation at 37°C for 30 minutes prior to flow cytometric analysis. After expansion for 4-5 days, the top 1% of HaloTag Ligand expressing cells were sorted as single cells into 96-well plates containing complete culture media and incubated at 37°C for expansion. LOUCY cells were difficult to expand from single clones, so we sorted cells into a 1:1 mixture of fresh media and 2-day conditioned media. After expansion, subclones were screened for homozygous dTAG-HaloTag insertion by PCR from crude lysates, then expanded further to confirm knock-in via Sanger sequencing and Linear Amplicon Nanopore sequencing (Eurofins Genomics).

### Validating LDB1-dTAG-HaloTag isogenic cell lines

LOUCY^DH^ and KOPT K1^DH^ subclones were first analyzed for similar levels of LDB1 expression to parental cells via Western blot. As LDB1 is a transcriptional coregulator and a bulky tag may influence this function, we next performed RNA-seq and ChIP-seq to ensure similar transcriptional and chromatin occupancy profiles compared with parental cell lines. To verify the efficacy of the dTAG system, we treated dTAG-HaloTag cell lines with 0-2,000 nM dTAG^V^-1 ligand^53^ (Tocris) for 0-24 hours. Loss of LDB1-dTAG-HaloTag expression was confirmed via Western blot and flow cytometry. All “acute” degradation experiments were performed with 500 nM dTAG^V^-1 or an equivalent volume of DMSO as a negative control for 4 hours for both cell lines; DMSO volume was normalized across all conditions at ≤0.1%. All experiments were performed in LOUCY^DH1^ and KOPT K1^DH1^ unless otherwise stated.

### CRISPR/Cas9 competitive cell proliferation assay

LOUCY, KOPT K1, JURKAT, SUP-T1 and PF-382 cell lines were transduced with a lentiviral vector encoding S.p. Cas9-P2A-PuroR^92^ (a gift from Christopher Vakoc [Addgene plasmid #108100]). Stable cell lines expressing the Cas9-Puro construct were selected by adding puromycin at 1 µg/mL (LOUCY) or 2 µg/mL (all other cell lines) for 5 days. Concentrations were based on kill curves in untransduced cells and resulting cell lines were maintained under puromycin selection at the same doses. The CRISPR Guide RNA Design Tool (https://benchling.com) was used to design 4 gRNAs targeting several LDB1 exons. Oligonucleotides were cloned into LRG2.1^92^ (a gift from Christopher Vakoc [Addgene plasmid # 108098]) and then transduced into Cas9-Puro T-ALL cell lines. NTC and PCNA-targeting gRNAs were used as negative and positive controls, respectively. At 72 hours post-transduction (Passage 0), GFP expression was checked by flow cytometry and confirmed to be 20-80%. 1e5 cells per condition were seeded into 96-well plates. 3-5e5 GFP-positive cells per LDB1-targeting gRNA were separately sorted to ensure loss of LDB1 protein expression via Western blot compared with untransduced controls and NTCs. Every 3 days, GFP percentage was measured using the FACSymphony High Throughput Sampler, and cells were passaged into fresh media. Final percentage of GFP+ cells (compared with live cells) was divided by initial GFP percentage (Passage 0) to calculate relative change from baseline. All gRNA sequences are reported in **Supplemental Table 13**.

### CRISPRa competitive cell proliferation assay

LOUCY^DH^ and KOPT K1^DH^ cell lines were transduced with a lentiviral vector encoding dCas9-VPR-P2A-mCherry^93^ (a gift from Anna Obenauf [Addgene plasmid #154193]) and sorted to isolate a population of 100% mCherry-expressing cells. Cells were re-sorted prior to any experiments if mCherry percentage drifted below 80%. CRISPick^94,95^ (Broad Institute) was used to design 5 gRNAs targeting putative LDB1-dependent genes using hg38, CRISPRa SpyoCas9 settings. We prioritized gRNAs located 50-200 bp upstream of the TSS and cloned them into the LRG2.1 backbone. In a pilot experiment, we transduced LOUCY^DH^ and KOPT K1^DH^ cell lines with the CRISPRa and gRNA constructs (4-5 gRNA per gene), depleted LDB1 with dTAG^V^-1 and analyzed target gene expression compared with DMSO-treated controls. We selected two gRNAs per gene with the target gene expression values closest to the DMSO control for subsequent experiments. For the competitive proliferation assay, LRG2.1 constructs were transduced into CRISPRa T-ALL cell lines. At 72 hours post-transduction, GFP expression (gRNA-positive cells) was checked by flow cytometry and confirmed to be 20-50%. 1e5 cells were seeded into 96-well plates in duplicate, with one replicate each treated with DMSO or dTAG^V^-1. Every 3 days, GFP percentage was measured using the FACSymphony High Throughput Sampler, and cells were passaged into fresh media containing either DMSO or dTAG^V^-1. Final percentage of live mCherry+ GFP+ cells (compared with total events) was divided by initial percentage of live mCherry+ GFP+ cells (Passage 0) to calculate relative change from baseline. In a separate experiment, we sorted 2e5 mCherry+ GFP+ cells, split them across two wells of a 96-well plate and treated each replicate with DMSO or dTAG^V^-1 for 72 hours. The 96-well plate was centrifuged at 500g for 5 minutes at 4°C, supernatant was aspirated, and cells were directly lysed in TRIzol Reagent (Invitrogen). RNA was then isolated using the Direct-Zol-96 RNA kit (Zymo Research) with on-column DNase I digestion prior to elution and analysis by RT-qPCR. All gRNA sequences are reported in **Supplemental Table 13**.

### Statin treatment and metabolite supplementation

Pitavastatin (HY-B0144A) and Rosuvastatin Calcium (HY-17504) were purchased from MedChemExpress, resuspended in DMSO, aliquoted and stored at −80°C. Individual aliquots were thawed, diluted into working stocks and stored at −20°C for short-term use. For LDB1 degradation experiments, LOUCY^DH^ and KOPT K1^DH^ cells were seeded into an opaque white flat-bottomed 96-well plate and treated with DMSO or 500 nM dTAG^V^-1 in combination with pitavastatin (0-50 µM) or rosuvastatin (0-400 µM) for 72 hours; DMSO volume was normalized across all conditions at 0.1%. For metabolite supplementation, these conditions were replicated with the addition of 200 µM mevalonate, 10 µM GGPP or 100 µM SyntheChol. For LDB1 knockout experiments, LOUCY, KOPT K1, JURKAT, SUP-T1 and PF-382 Cas9-Puro cell lines were transduced with LDB1-targeting or non-targeting LRG2.1 constructs. At 48 hours post-transduction, we confirmed >80% GFP expression by flow cytometry, seeded cells into an opaque white flat-bottomed 96-well plate and treated them with pitavastatin (0-50 µM) or rosuvastatin (0-400 µM) for 72 hours; DMSO volume was normalized across all conditions at 0.1%. For all dose-response experiments, 2.5e4 cells were seeded in 50 µL of media (including drug/metabolite treatment) per well and performed as 3 biological replicates with technical duplicates each. After 72 hours of drug/metabolite treatment, cells were lysed with the CellTiter-Glo 2.0 reagent (Promega) per manufacturer specifications and luminescence was measured with a Tecan Infinite M200 Pro Plate Reader. Relative luminescence was normalized to untreated (no statin) wells for each combination, and dose-response curves were fit non-linearly with a variable slope (4 parameters) in GraphPad Prism v10.2.2.

### Fatostatin treatment

Fatostatin (S9785) was purchased from SelleckChem, resuspended in DMSO, aliquoted and stored at −80°C. Individual aliquots were thawed, diluted into working stocks and stored at −20°C for short-term use. We performed an initial pilot experiment to test transcriptional downregulation of SREBP2-target genes using 0, 6, 12, 24 or 48 hours of fatostatin treatment and observed that 24 hours was sufficient to elicit transcriptional changes without compromising viability. For subsequent experiments, LOUCY^DH^ and KOPT K1^DH^ cells were pre-treated with fatostatin for 20 hours. 500 nM dTAG^V^-1 or DMSO was then directly added to fatostatin-treated or untreated cells for an additional 4 hours prior to harvest, RNA extraction and RT-qPCR. KOPT K1^DH^ cells were treated with 0, 5, 10 or 20 µM fatostatin. LOUCY^DH^ cells were treated with 0, 5, or 10 µM fatostatin because the 20 µM fatostatin/dTAG^V^-1 combination resulted in near-complete cell death at 24 hours.

### ChIP-seq analysis

A detailed protocol for ChIP-seq library preparation can be found in the **Supplemental Methods**. All antibodies for ChIP-seq are reported in **Supplemental Table 14**. Libraries were sequenced in paired-end mode on an Illumina Novaseq X Plus platform at Novogene (2 x 150 bp reads) or on an Illumina NextSeq 2000 platform (2 x 50 bp reads).

ChIP-seq was performed for 2-4 biological replicates for each cell line, antibody and treatment. Input material corresponding to each cell line and treatment were also sequenced. Reads were aligned to the hg38 reference genome using Bowtie2^96^ v2.5.1. Reads with MAPQ <20 were filtered out using SAMtools^97^ v1.17. Duplicate reads were removed using Picard v3.0.0 (https://broadinstitute.github.io/picard/). We first generated bigwig files for each replicate using deeptools^98^ v3.5.1 bamCoverage. Concordance across replicates for treatments/antibodies was confirmed using multiBigwigSummary and plotCorrelation to calculate Pearson correlations. We then merged BAM files using SAMtools merge and made summary bigwig files using bamCoverage with unified hg38 blacklist region^99^ subtraction (ENCODE file <u>ENCFF356LFX</u>) with the following parameters: --binSize 20 --smoothLength 60 --normalizeUsing BPM --ignoreDuplicates --centerReads --minMappingQuality 20 --ignoreForNormalization chrX chrY chrM. Peaks were called from merged BAM files using MACS2^100^ v2.2.7. callpeak using merged inputs from each cell line and each treatment (DMSO or dTAG^V^-1) as control files with the following parameters: -f BAMPE -g hs -q 0.01. Quality of ChIP-seq was assessed by calculation of Fraction of Reads in Peaks^101^ (FRiP). For SMC3 ChIP-seq, peaks within 1 kb were stitched together using BEDtools^102^ v2.31. merge. For enhancer and promoter annotations, a list of transcription start sites (TSSs) was generated from the hg38 RefSeq gene list. We then calculated H3K27ac peak centers and measured the distance to the nearest TSS using BEDtools closest. Merged H3K27ac peaks within 1 kb were annotated as active promoters, and those outside of 1 kb were annotated as active enhancers. Promoter regions within 1 kb of each other were stitched together, as were enhancer regions.

LDB1 peaks overlapping with enhancers or promoters, as well as overlaps between LOUCY and KOPT K1, were calculated using BEDtools intersect. Bigwig signal matrices for each ChIP were generated with deeptools computeMatrix using total LDB1 peaks or LDB1 peaks at enhancer regions in DMSO-treated conditions as reference files with unified hg38 blacklist region subtraction and the following parameters: --referencePoint center -b 3000 -a 3000 --missingDataAsZero. Heatmaps and profiles were then generated using deeptools plotHeatmap with the following parameters: --refPointLabel center --sortRegions descend --sortUsing mean. UpSet plots were generated with UpSetPlot^103^ v0.9.0; overlaps were defined based on the DMSO-treated conditions for each ChIP. HOMER^104^ v4.11. was used for motif analysis. Briefly,

HOMER findMotifsGenome.pl was used to calculate de novo and known motif enrichment within 200 bp of the center of LDB1-occupied peaks at enhancers that were unique to LOUCY, unique to KOPT K1 or shared between both cell lines. A union set of all enhancers with or without LDB1 occupancy was used as background for analysis. Next, using known motif enrichment, we classified redundant motifs based on transcription factor family (e.g. ETS, RUNX, GATA, bHLH, etc.) to create a heatmap of individual motif enrichment color-coded by the transcription factor family.

### TT-seq data analysis

TT-seq was performed as previously described^21,54,55,81^ with minor modifications. A detailed protocol for TT-seq library preparation can be found in the **Supplemental Methods**. Libraries were pooled and sequenced on the Illumina NextSeq 2000 platform to generate 2 x 50 bp reads using Illumina sequencing reagents according to the manufacturer specifications.

TT-seq analysis was performed as previously described^21,55,81,105^. The paired-end TT-seq reads were trimmed using Trim Galore v0.6.10 (https://github.com/FelixKrueger/TrimGalore/) and were aligned to the human hg38 and Drosophila dm6 assemblies and sorted using STAR v2.7.10b. Index, skip, MAPQ <7 alignments in the resulting BAM files were filtered out using SAMtools v1.14. Duplicate reads were removed with Picard v3.0.0 MarkDuplicates. Raw read counts across gene bodies were quantified using HOMER v4.11.1. Once replicate correlation was confirmed by principal component analysis, we merged unnormalized BAM files across replicates for each cell line and treatment using SAMtools v1.17 merge and generated bigwig files using deeptools v3.5. bamCoverage with CPM normalization. Merged bigwigs were then used for visualization with Micro-C contact maps. We compared differential expression analysis between spike-in based scale factors and default DESeq2 settings and observed high concordance. Therefore, DESeq2 default normalization was used for our final analysis using variance-stabilizing transformation (VST) and a significance threshold of Benjamini-Hochberg corrected *P_adj_* <0.05. The *apeglm* method was used for log_2_-fold change shrinkage. Batch correction was performed for KOPT K1 samples as these were run on two flowcells. We used Gene Set Enrichment Analysis^106,107^ v4.4.0 to calculate enrichment scores from TT-seq datasets by ranking all genes for either cell line by the DESeq2 Wald statistic (“stat”), then applying the GSEAPreranked algorithm with 1,000 permutations using the v2025 Human Collection MSigDB.

### Micro-C data processing

Micro-C was performed as previously described^21,22,56–58,81,108^ with several modifications. A detailed protocol for Micro-C library preparation can be found in the **Supplemental Methods**. Shallow sequencing libraries were prepared in paired-end mode on the Illumina NextSeq 2000 platform to generate 2 x 50 bp reads. Deep sequencing of high-quality libraries was then performed in paired-end mode on the Illumina Novaseq X Plus platform at Novogene to generate 2 x 150 bp reads.

Micro-C analysis was performed as previously described^21,22,81^ (https://github.com/jclqrs/Lam_2024_Code). We used the distiller pipeline v3.3 (https://github.com/open2c/distiller-nf/) to generate contact maps using FASTQ files as input. PCR duplicates were removed from each replicate, and balanced contact maps were generated for each treatment from merged biological replicates. Iterative correction and eigenvector decomposition (ICE) balancing was used to normalize contact maps using default settings: a given bin was excluded if its sum was >5 median absolute deviations below the median bin, the first two diagonals were ignored for balancing, columns and rows were normalized so that they summed to 1. We used CoolBox^109^ v0.3.8 to visualize contact maps and aligned ChIP-seq and TT-seq tracks. To generate aggregate pileup plots (APA plots) of Micro-C contacts over regions (defined in figure legends) and contact-decay P(s) curves, we used cooltools^110^ v0.7.1.

Micro-C compartment analysis, domain analysis, and loop calling and quantification were performed as previously described with minor modifications^21,22,81,83^ using cooltools v0.7.1, rGMAP^111^ v1.4 and cooltools v0.7.1, respectively. A detailed protocol can be found in the **Supplemental Methods**.

### Loop classification

To classify loops, we used CTCF, SMC3 and LDB1 ChIP-seq peaks, as well as active enhancer/promoter annotations based on H3K27ac ChIP-seq peak distance from TSSs as previously described. We normalized loop anchors to 10 kb by taking the center point of each anchor and extending it 5 kb in either direction. We used BEDtools v2.31.1 intersect to characterize whether 0, 1 or 2 loop anchors overlapped (≥1 bp) with LDB1 peaks, active enhancer regions/promoter regions or CTCF/SMC3 co-occupied peaks. We then classified loops as 1) structural loops (CTCF/SMC3 co-occupancy at both anchors with enhancer and/or promoter occupancy at 0 or 1 anchor), 2) dual-function loops (CTCF/SMC3 co-occupancy at both anchors with enhancers and/or promoters at both anchors), 3) CRE loops (enhancers and/or promoters at both anchors and CTCF/SMC3 co-occupancy at 0 or 1 anchor) or 4) unclassified (not structural, dual-function or CRE). We next categorized CRE loops and dual-function loops as enhancer-enhancer loops (enhancers at both anchors without promoters), enhancer-promoter loops (enhancer at one anchor, promoter at the opposite anchor), promoter-promoter loops (promoters at both anchors without enhancers) or mixed loops (enhancers and promoters were within the same anchor). We then stratified structural, dual-function and CRE loops based on whether LDB1 peaks were present at 0, 1 or 2 anchors. LDB1-dependent loops were defined as loops with LDB1 peaks at 1 or 2 anchors that were weakened upon LDB1 depletion. LDB1-independent loops were defined as loops that were unchanged/strengthened upon LDB1 depletion or weakened without LDB1 at either anchor.

Loop calling was also performed using mustache^112^ (merging 10, 5, and 2 kb resolution loop calls) and peakachu^113^ (merging 10 and 5 kb resolution loop calls) with quantification of loop strength using cooltools. A similar trend was observed in which a higher percentage of CRE loops were weakened upon LDB1 depletion than structural or dual-function loops. Our final analysis uses cooltools for consistency between loop calling and loop strength calculation.

### Integration of looping with transcription

To integrate looping with transcription, we defined a 5 kb window around TSSs of active genes from our TT-seq data. We then intersected loop anchors with the TSS windows for LDB1-dependent and LDB1-independent CRE or dual-function loops. Genes were further stratified by the number of loop anchors overlapping with the TSS window. We then calculated gene expression changes based on the DESeq2 Wald statistic from our TT-seq data, which takes into account both fold change as well as statistical significance. Data in **Fig. 3e** depict LDB1-dependent genes; analysis of all genes confirmed a statistically significant association between gene expression and LDB1-dependent loops with an expected decrease in effect size due to a high proportion of unchanged genes.

### Loop rewiring analysis

Genuine anchor-level rewiring was defined for each weakened CRE loop anchor that overlapped a strengthened loop anchor, subject to the requirement that the new loop partner (the other anchor of the strengthened loop) did not overlap the original loop partner (the other anchor of the weakened loop), ensuring that only bona fide partner switching, rather than loop expansion or contraction, was counted. Weakened CRE loops were stratified by LDB1 occupancy at their anchors (LDB1 at 0 or ≥ 1 anchors), and the fraction of loops undergoing genuine rewiring was compared between groups using Fisher’s exact test in LOUCY^DH^ and KOPT K1^DH^ cells independently. For each rewired weakened CRE loop, the strengthened partner loop was classified as CRE, dual-function, both or unclassified (non-structural), and the distribution of rewiring destinations was visualized per unique weakened loop stratified by LDB1 status. To assess the transcriptional consequences of rewiring, genes within 5 kb of each anchor were linked to TT-seq results and stratified into downregulated, not significant or upregulated categories, allowing assessment of whether rewired contacts were preferentially associated with transcriptionally upregulated genes.

### HiChIP data processing and comparison to Micro-C

H3K27ac HiChIP data from primary T-ALL patient samples were downloaded from previous publications^36,47^. For analysis of HiChIP overlap with Micro-C data, raw reads were aligned to the hg38 reference genome using the juicer pipeline^114^ v2.20.00. To identify H3K27ac peaks from each sample, deduplicated alignments with insert size <1 Mb were first selected using SAMtools v1.9 and converted to BED format using BEDtools v2.30.0. Peaks were called from the filtered alignments using MACS2 with parameters: -g hs -q 0.05. As cooltools uses local background to call loops, this limited our analysis of HiChIP data due to areas of sparse read depth. We therefore used FitHiChIP^115^ to call HiChIP loops with parameters: CircularGenome=0, IntType=3, BINSIZE=10000, LowDistThr=20000, UppDistThr=2000000, UseP2PBackgrnd=0, BiasType=1, MergeInt=1, QVALUE=0.01, which takes as input alignments with mapping quality score greater than 30 (inter_30.hic) and H3K27ac peaks. Significant interactions (contacts) with H3K27ac peaks detected at both anchors (representing dual-function or CRE loops) were overlapped with Micro-C CRE and/or dual-function loops defined in LOUCY or KOPT K1 using BEDtools pairtopair with the requirement that both loop anchors overlapped. HiChIP datasets with fewer than 10,000 H3K27ac peaks were filtered out. Micro-C loops that overlapped with >1 HiChIP loop were deduplicated as we could not distinguish whether these represented biologically distinct loops or artifactual differences from different techniques. We then calculated the ratio of overlap between ETP/near-ETP ALL and non-ETP ALL HiChIP samples to LOUCY or KOPT K1 Micro-C.

For loop strength analysis, we used the distiller pipeline v3.3 to generate mcool files for each H3K27ac HiChIP file using FASTQ files as input with the same workflow described for our Micro-C data. We used CoolBox v0.3.8 to visualize and compare HiChIP and Micro-C contact maps with aligned ChIP-seq and TT-seq tracks. We generated aggregate pileup plots (APA plots) for each HiChIP dataset at 10 kb resolution over all CRE loops, LDB1-dependent CRE loops and LDB1-dependent dual-function loops using cooltools v0.7.1. The log_2_(observed/expected) score was then used to compare HiChIP loop strength at contacts defined from LOUCY or KOPT K1 Micro-C experiments, and each point was color coded based on their differentiation state (ETP/near-ETP, non-ETP or normal controls) based on annotations from the original publications.

### Integration of TT-seq with primary T-ALL bulk and single-cell transcriptomics

Differentially expressed genes were obtained from TT-seq experiments in KOPT K1 and LOUCY cell lines. For each cell line, genes were partitioned into upregulated and downregulated sets based on a Benjamini-Hochberg corrected *P_adj_* <0.05 and log_2_(FC) >|0.5|. Gene expression for primary samples (<u>syn54032669</u>)^36^ was represented as a VST matrix (DESeq2 v1.38.3), with genes in rows and samples in columns. Per sample gene set activity scores were computed using GSVA^116^, which estimates an enrichment-like score per gene set and sample in an unsupervised manner. GSVA tool v2.0.0 was run on the VST expression matrix using the GSVA package default settings. For visualization and survival analyses, gene set scores were Z-score standardized across samples. Event-free survival and overall survival times and event indicators were obtained from curated clinical annotations^36^. Patients with missing time or event status were excluded per endpoint. For each gene set, patients were stratified into tertiles based on the standardized continuous gene set score, and Kaplan Meier curves were generated for the three groups using survival R package v3.8-3. Group differences were assessed using the log-rank test.

Univariable models were fit separately for each GSVA score. Multivariable models were fit to evaluate whether GSVA scores retained prognostic significance after adjustment for clinical covariates, minimal residual disease, white blood cell count, central nervous system status, and sex. Minimal residual disease was modeled as log(MRD fraction + 1e-5). Hazard ratios and 95% confidence intervals were reported per one standard deviation increase in GSVA score. *P*-values were adjusted for multiple testing within each endpoint and model using the Benjamini-Hochberg false discovery rate procedure. Analyses were performed in the full cohort and in subtype restricted subsets, including ETP-like and TAL1 related samples.

For comparison of TT-seq upregulated/downregulated gene sets to single-cell RNA-seq (scRNA-seq) datasets^60,61^, signature scores were calculated from log-normalized expression using the scanpy function tl.score_genes using healthy pediatric hematopoiesis (49,623 cells) and T-ALL (114,817 cells) reference datasets. Sub-analysis was also performed for LOUCY gene set expression in an ETP ALL reference dataset (69,483 cells). T-cell developmental arrest phenotypes were stratified as BMP (HSPC through ETP) versus Late T (Pro-T through Effector T) or BMP (HSPC through ETP) versus pre-positive selection (Pro-T through DP) versus post-positive selection (alpha-beta through Effector T).

### Statistics

No statistical method was used to predetermine sample size. All statistical details for experiments can be found in the figure legends. GraphPad Prism v10.2.2 was used to calculate one- or two-way ANOVA tests, unpaired Student’s *t*-tests or Fisher’s exact tests to compare categorical variables and the extra sum-of-squares F test for dose-response curves. The SciPy v1.15.2 package in Python v3.10.16 was used to calculate Kruskal-Wallis tests or Dunnett’s test (two-sided, FWER <0.05). Multiple comparison testing as appropriate was performed and described in figure legends. Unless otherwise noted, data are summarized as means ± standard deviation. *P*-values or *q*-values <0.05 were considered significant and are reported in figures or denoted by * (*P* or *q* < 0.05), ** (*P* or *q* < 0.01), *** (*P* or *q* < 0.001), **** (*P* or *q* <0.0001) or ns (*P* or *q* ≥ 0.05).

## Data availability

RNA-seq, ChIP-seq, ATAC-seq, Micro-C and TT-seq data generated in this study will be publicly available in the NCBI Gene Expression Omnibus (GEO) at the time of publication. Data sources from previously published studies can be found at: H3K27ac HiChIP datasets (GEO Accession <u>GSE243915</u>, European Genome-phenome Archive Accession <u>EGAS50000000016</u>), bulk RNA-seq from primary T-ALL samples (dbGaP accessions <u>phs002276</u> and <u>phs000464</u>, Synapse accession <u>syn54032669</u>), scRNA-seq from primary T-ALL samples and healthy pediatric hematopoiesis controls (dbGaP accession <u>phs003432</u>, <u>Human Tumor Atlas Network</u>), RNA-seq from *Lmo2* transgenic mouse thymocytes (GEO accession <u>GSE129244</u>); RNA-seq from human post-natal thymocytes (GEO accession <u>GSE151079</u>). Original western blot images are included as a **Supplemental** attachment. Any additional data required to reanalyze the data in this study is available upon request to the lead contacts.

## Code availability

This manuscript does not report original code. Previously published code for Micro-C analysis is available at https://github.com/jclqrs/Lam_2024_Code. Any additional code required to reanalyze the data in this study is available upon request to the lead contacts.

## Acknowledgements

We would like to thank the Blobel Lab members for their discussions; mentors from the Translational Research Training in Hematology Program; Dr. Wei Tong for sharing mTOR/AKT/S6 antibodies used in this study; Dr. Theresa Palomero for providing the CUTLL3 cell line; Dr. Robert Babak Faryabi for providing the KOPT K1 cell line; the Children’s Hospital of Philadelphia Flow Cytometry Core for assistance; Huck Institutes’ Genomics Research Incubator (RRID:SCR_024530) for use of the Illumina NextSeq 2000. Schematics in **Figs. 1, 4**, and **5** were created with <u>BioRender.com</u>.

## Author contributions

R.S.B. and G.A.B. conceived the study and designed the experiments. R.S.B. engineered the LDB1-dTAG-HaloTag cell lines and performed RNA-seq, ATAC-seq, ChIP-seq, Micro-C and TT-seq experiments and analyzed the data. S.W., S.Z., Z.G. and M.C.A. assisted with data processing for Micro-C, HiChIP, ChIP-seq and TT-seq datasets and assisted with data analysis. J.S.L. and A.T. assisted with CRISPR, Western blot, RT-qPCR and cell viability experiments in this study. P.P. and C.G.M. provided raw HiChIP data from primary T-ALL samples and performed comparative analysis of TT-seq datasets to primary sample RNA-seq datasets, including prognostic analyses. T.I. and K.T. performed comparative analysis of TT-seq datasets to primary sample scRNA-seq datasets. ATAC-seq and RNA-seq library preparation and next-generation sequencing were performed by C.A.K. and B.M.G. under the supervision of R.G.H. N.G.A. and S.J.S. provided critical insights. The manuscript was written by R.S.B. and G.A.B. and reviewed and edited by all authors.

## Ethics Declarations

### Materials and Correspondence

Correspondence to:

Rahul S Bhansali

Abramson Research Center 315 3615 Civic Center Blvd.

Philadelphia, PA 19104-4318

Phone: (630) 303-7306

Gerd A Blobel, MD, PhD Abramson Research Center 316H 3615 Civic Center Blvd.

Philadelphia, PA 19104-4318

Phone: (215) 590-3988

