## Supplemental Methods and Figs. S1-S9 for "LDB1-dependent enhancer connectivity defines T-cell leukemia identities and masks metabolic vulnerabilities"

**Supplemental Appendix:**

**I. Supplemental Methods**

**II. Supplemental References**

**III. Supplemental Data:**

**Supplemental Fig. S1.** Orthogonal validation of LDB1 knockout and dTAG systems, related to **Fig. 1**.

**Supplemental Fig. S2.** Representative flow cytometry scatterplots, related to **Fig. 1**

**Supplemental Fig. S3.** Subtype-specific transcription factors govern LDB1/LMO2 complex recruitment to enhancers, related to **Fig. 2.**

**Supplemental Fig. S4.** Acute LDB1 depletion disrupts T-ALL subtype-specific oncogenic transcriptional programs but minimally impacts larger scale chromatin structure, related to **Fig. 3.**

**Supplemental Fig. S5.** Acute LDB1 depletion preferentially disrupts cis-regulatory loops, related to **Fig. 3.**

**Supplemental Data Figure S6.** Acute LDB1 depletion promotes rewiring of cis-regulatory element loops, related to **Fig. 3.**

**Supplemental Fig. S7.** LDB1 drives T-ALL subtype identities and constrains against alternative transcriptional states, related to **Fig. 5.**

**Supplemental Fig. S8.** Representative Micro-C contact maps for mevalonate pathway gene chromatin loop rewiring, related to **Fig. 7.**

**Supplemental Fig. S9.** Mevalonate pathway phenotypic changes induced by LDB1 loss are conserved in human T-ALL cell lines and mouse models of T-ALL, related to **Fig. 7.**

**IV. Supplemental Tables 1-14:**

**Supplemental Table 1.** List of human T-ALL cell lines and notable genetic features

**Supplemental Table 2.** HOMER motif enrichment under LDB1 peaks at enhancers for LOUCY$^{DH1}$ and KOPT K1$^{DH1}$

**Supplemental Table 3.** LOUCY$^{DH1}$ TT-seq DESeq2 table

**Supplemental Table 4.** KOPT K1$^{DH1}$ TT-seq DESeq2 table

**Supplemental Table 5.** LOUCY$^{DH1}$ RNA-seq DESeq2 table

**Supplemental Table 6.** KOPT K1$^{DH1}$ RNA-seq DESeq2 table

**Supplemental Table 7.** Micro-C quality metrics

**Supplemental Table 8.** LOUCY$^{DH1}$ Micro-C loop calls with classification

**Supplemental Table 9.** KOPT K1$^{DH1}$ Micro-C loop calls with classification

**Supplemental Table 10.** HiChIP quality metrics and overlap with Micro-C

**Supplemental Table 11.** Raw data for GSVA univariable/multivariable analysis

**Supplemental Table 12.** Correlation of LDB1-dependent transcriptional profiles (GSVA from TT-seq) with genetic lesions via two-sided Wilcoxon test

**Supplemental Table 13.** List of oligonucleotides for cloning, qPCR and gRNA synthesis

**Supplemental Table 14.** List of antibodies used for flow cytometry, Western blots and ChIP

#### Supplemental Methods

##### T-ALL cell lines

Human T-ALL cell lines were purchased from The Leibniz Institute DSMZ (DND-41, MOLT-14, PEER, P12 ICHIKAWA, PF-382, SUP-T1), ATCC (LOUCY, JURKAT Clone E6-1), or Takara Bio (Lenti-X 293T). The CUTLL3 and KOPT K1 human T-ALL cell lines were gifted by Dr. Theresa Palomero and Dr. Robert Babak Faryabi, respectively. Upon receipt, cell lines were thawed, expanded, confirmed to be free of mycoplasma contamination by MycoAlert (Cambrex) through the University of Pennsylvania Department of Genetics Cell Center (Philadelphia, PA), and authenticated by a 9-marker STR profile test through the Penn Genomics and Sequencing Core Facility (Philadelphia, PA). Mycoplasma testing and STR authentication were repeated before any major experiments and before cryopreservation for all parental and derivative cell lines. LOUCY, KOPT K1, CUTLL3, and PEER were grown in RPMI (Gibco or Cytiva) supplemented with 20% FBS (HyClone), 2 mM GlutaMAX (Gibco), 1 mM Sodium Pyruvate, 1x MEM Non-Essential Amino Acid Solution (Gibco), 55 µM beta-mercaptoethanol (Gibco), and Normocin (Invivogen). All other T-ALL cell lines were grown in RPMI supplemented with 10% FBS, 2 mM GlutaMAX, and Normocin. Cell lines were passaged every 2-3 days to maintain a density of 7.5e5-1e6 cells/mL (LOUCY and PEER) or 3-5e5 cells/mL (all other cell lines).

##### Primary tissue samples

Frozen CD34+ hematopoietic stem and progenitor cells (HSPCs) purified from peripheral blood mononuclear cells of single donors were obtained from the Fred Hutchinson Cooperative Center for Excellence in Hematology (Seattle, WA). Cells were thawed at day 0 according to core-supplied protocol and cultured using a previously described protocol[1]. Briefly, HSPCs were expanded for 5 days in StemSpan SFEM-II (STEMCELL Technologies) supplemented with 100 ng/mL SCF, 100 ng/mL FLT3L, 100 ng/mL TPO, 20 ng/mL IL-6, and 50 ng/mL IL-3. For proerythroblast differentiation, CD34+ HSPCs were differentiated for 6 days in StemSpan SFEM-II supplemented with 50 ng/mL SCF, 3 U/mL erythropoietin, 20 ng/mL IL-3, and 40 ng/mL IGF-1.

For primary human thymic samples, subjects were consented and enrolled in the CHOP Birth Defects Biorepository (BDB) between 2023 and 2024 under an Institutional Review Board approved protocol (IRB 18–015525). Abstracted and deidentified data from the electronic health record were provided by the BDB to study investigators, which included demographics, clinical data, and clinical genetic testing results. De-identified thymus tissue collected from these enrolled subjects and stored in the Birth Defects Biorepository was also provided. Thymi were resected in the cardiac operating room as a necessity for certain cardiac lesion repairs. They were then placed on ice, transported to the laboratory, and then aliquoted and stored. Paired flash frozen tissue or tissue stored in RNA*later* (Invitrogen) were provided in 50mg aliquots.

**DepMap analysis**

We used the Custom Analysis tool in the DepMap Portal[2] (Broad Institute, DepMap Public 25Q2 release) to compute Pearson correlation between Chronos Gene Effect and LMO2 gene expression. Scatter plot is filtered on cancer cell lines of hematopoietic lineage (OncotreeLineage Lymphoid and Myeloid).

**Lentiviral packaging and transduction**

All lentiviral plasmids were packaged using Lenti-X 293T cells (Takara Bio) at low passage (3-4 passages). Briefly, Lenti-X 293T cells were grown to 70-90% confluency. At least 1 hour prior to transfection, we exchanged cells into fresh media and equilibrated them at 37°C. For lentiviral packaging, plasmid DNA was mixed with psPAX2 and VSV-G (2:1.5:1 molar ratio). Polyethyleneimine (Millipore Sigma) was then added to the plasmid mixture at a ratio of 4:1 (w/w) in Opti-MEM I (Gibco). After 18 minutes of room temperature incubation, the transfection cocktail was added dropwise to 293T cells and incubated at 37°C prior to exchanging media after 4-8 hours. Viral supernatant was harvested at 24, 48, and 72 hours after transfection, filtered using 0.45 µm polyethersulfone (Millipore Sigma) or cellulose acetate (Fisher Scientific) filters, concentrated 100-fold using Lenti-X Concentrator (Takara), and stored at -80°C in single use aliquots for up to 6 months. For transduction, T-ALL cell lines were seeded into 24-well plates at 5e5-1e6 cells/mL prior to adding concentrated lentivirus, 25 mM HEPES, and either 1:50 LentiBoost Research Grade Solutions A and B (Mayflower Bioscience) for LOUCY and KOPT K1 or 8 µg/mL polybrene for JURKAT, PF-382, and SUP-T1. Cells were spinoculated at 800g x 90 minutes at room temperature and incubated overnight prior to washing and resuspending into fresh media.

**Flow cytometry and cell sorting**

Flow cytometric analysis was performed on a FACSymphony or LSR Fortessa Flow Cytometer (BD) and analyzed with FlowJo v10. Cell sorting was performed on a FACSAria or FACSJazz. For analysis of fluorescent markers (GFP, mCherry, or HaloTag), cells were centrifuged at 500g x 5 minutes at 4°C, washed twice in cold 1x PBS, resuspended in FACS Buffer (1x PBS supplemented with 2% FBS and 2 mM EDTA) prior to analysis. For cell cycle/senescence analysis, 10 µg/mL Hoechst 33342 (Invitrogen) was added directly to 5e5 cells in culture with 1 hour incubation at 37°C. Cells were then centrifuged at 500g x 5 minutes at 4°C, washed twice in cold 1x PBS, and resuspended in 100 µL FACS buffer supplemented with 1 µg/mL pyronin Y (Millipore Sigma) followed by 15 minutes incubation at room temperature protected from light. The CellEvent Senescence Green Flow Cytometry Assay Kit (Invitrogen) was then used to fix and stain for β-galactosidase per manufacturer specifications. For cell death analysis, cells were centrifuged at 500g x 5 minutes at 4°C, washed twice in cold 1x PBS, resuspended in Annexin Binding Buffer (BD) at 1e6 cells/mL, and supplemented with 5 µL each of Annexin V-FITC (BD) and Propidium Iodide (BD). Cell suspensions were incubated for 15 minutes at room temperature prior to dilution with Annexin Binding Buffer and analysis. For all experiments, unstained and single-color controls were used for compensation, and cells were passed through a 40 µm mesh filter prior to analysis.

**Western blotting**

T-ALL cell lines, primary CD34+ HSPCs, and primary proerythroblast cells were centrifuged at 500g x 5 minutes at 4°C and washed twice with cold 1x PBS. Pellets were then resuspended in RIPA Buffer (50 mM Tris-HCl pH 8, 150 mM NaCl, 1% NP-40, 0.5% sodium deoxycholate, 0.1% SDS) supplemented with 1x EDTA-free cOmplete protease inhibitor cocktail (Roche), 1 mM PMSF, 2 mM NaF, 2 mM $NaVO_3$, 2 mM beta-glycerophosphate, 2 mM MgCl2, and 250 U/mL Benzonase (Millipore Sigma) and incubated at 4°C with intermittent agitation for 30 minutes. Lysates were then sonicated with the Bioruptor Pico (Diagenode) for 10 minutes (30 seconds on/off, "easy" mode). Lysates were clarified by centrifugation at 15,000g for 10 minutes at 4°C. Frozen thymic samples (50mg) were homogenized with 50mg Protein Extraction Beads (Diagenode) in 250µl supplemented RIPA buffer. Samples were agitated at 4°C for 20 minutes, sonicated for 10 cycles (30 seconds on/off, “easy” mode), and agitated again at 4°C for 20 minutes. Supernatants were transferred to new tubes, leaving beads behind, and then clarified by centrifugation.

All protein lysates were quantified with the Pierce 600 nm Protein Assay (Thermo Scientific) and denatured with 4x Sample Buffer (LI-COR) at 95°C for 5 minutes. Equal amounts of protein lysates (5-15 μg) were loaded onto 4-12% NuPAGE Bis-Tris Mini or Midi protein gels and run with NuPAGE MOPS SDS Running Buffer. Proteins were transferred onto low-fluorescence PVDF membranes, dried, re-activated with methanol, and stained for total protein using Revert 700 Total Protein Stain (LI-COR). After reversal of total protein stain, membranes were blocked for 1 hour at room temperature with Intercept Blocking Buffer (LI-COR), followed by overnight incubation with primary antibodies at 4°C diluted in Intercept T20 Antibody Diluent (LI-COR). Membranes were then washed with TBS-T, incubated with IRDye secondary antibodies (LI-COR) for 1 hour at room temperature, and washed again with TBS-T. Total protein stain and fluorescent antibody signals were detected using the LI-COR Odyssey imaging system. All antibodies are reported in **Supplemental Table 14**.

**Reverse Transcriptase Quantitative Polymerase Chain Reaction**

RNA from T-ALL cell lines, primary CD34+ HSPCs, and primary proerythroblast cells was extracted using the RNeasy Plus Mini Kit (Qiagen). Cells were centrifuged at 500g x 5 minutes at 4°C and immediately resuspended in RLT Plus Buffer supplemented with beta-mercaptoethanol. Samples were homogenized using QIAshredder columns, and genomic DNA was removed using gDNA Eliminator Columns and on-column DNase digestion. For primary thymic tissue, excess

RNA*later* was blotted off and then added to a dounce homogenizer with 1 mL TRIzol Reagent (Invitrogen). Tissue was homogenized with 30 passes, centrifuged at 12,000g x 5 minutes at 4°C to remove excess fatty tissue, and supernatant was transferred to a new tube for TRIzol Reagent RNA extraction per manufacturer specifications. Purified RNA was then diluted in RLT Plus Buffer supplemented with beta-mercaptoethanol and re-purified using the RNeasy Plus Mini Kit (Qiagen) to allow for on-column DNase digestion. RNA was quantified by NanoDrop (Thermo Scientific), and cDNA was synthesized using iScript Reverse Transcription Supermix (Bio Rad). "No RT" negative controls were used for primary transcript detection to rule out genomic DNA contaminants. cDNA was diluted 5- or 10-fold, and qPCR was performed using Power Sybr Green PCR Master Mix (Applied Biosystems) on either Viia 7 or QuantStudio 5 platforms. For detection of mRNA, normalization was performed against the geometric mean of Ct values for *ACTB, GAPDH, B2M,* and *ABL1*. For detection of primary transcripts, normalization was performed against the geometric mean of Ct values for *ACTB, GAPDH, TOP2A,* and *RPL35*. Relative expression was calculated using the $2^{-\Delta\Delta Ct}$ method. All primers are reported in **Supplemental Table 13**.

**RNA-seq and data analysis**

Total RNA was extracted from parental LOUCY and KOPT K1 cells, as well as LDB1-dTAG-HaloTag clones using the RNeasy Plus Mini Kit (Qiagen). Cells were centrifuged at 500g x 5 minutes at 4°C and immediately resuspended in RLT Plus Buffer supplemented with beta-mercaptoethanol. Samples were homogenized using QIAshredder columns, and genomic DNA was removed using gDNA Eliminator Columns and on-column DNase digestion. RNA Integrity Number (RIN) was confirmed to be 8.9-9.7 for all samples using the Bioanalyzer RNA 6000 Nano

(Agilent). RNA-sequencing libraries were prepared using 500 ng of total RNA using the Illumina Stranded mRNA Library Prep Kit (Illumina) according to manufacturer's specifications. Specifically, polyA+ selection was performed, followed by fragmentation and reverse transcription to produce cDNA. Sequencing adapters were then added via ligation, and unique dual indexes were added to the fragments via PCR. Following 10 cycles of PCR amplification, any remaining free adapters were removed via a 1x bead clean up using AMPureXP beads (Beckman Coulter).

The size and quality of each library were then evaluated by Bioanalyzer 2100 (Agilent Technologies) and quantified using qPCR (New England Biolabs). Libraries were sequenced in paired-end mode on the NextSeq 2000 platform to generate 2 x 50 bp reads using Illumina-supplied kits as appropriate. The sequence reads were processed using the ENCODE3 long RNA-seq pipeline (https://www.encodeproject.org/pipelines/ENCPL002LPE/). Briefly, reads were mapped to the human genome (hg38 assembly) using STAR[3], followed by RSEM[4] for gene quantifications. Differential expression analysis was performed with the DESeq2[5] v1.42 R package using Variance Stabilizing Transformation (VST) and a significance threshold of Benjamini-Hochberg corrected $P_{adj} < 0.05$. The *apeglm* method[6] was used for $\log_2$-fold change shrinkage.

**OMNI-ATAC-seq and data analysis**

Live LOUCY$^{DH}$ and KOPT K1$^{DH}$ cells were isolated using the MACS Dead Cell Removal Kit (Miltenyi Biotec) and resuspended in fresh media. The following morning, we confirmed >90% viability via Trypan blue staining prior to DMSO or dTAG$^{V}$-1 treatment. For each sample, 1e5 cells were collected in Protein LoBind tubes (Eppendorf) and centrifuged at 300g for 5 minutes at

4°C, after which the supernatant was discarded. Cells were washed with 1 mL of cold 1x PBS, spun as above, and the supernatant was discarded. We then proceeded with the OMNI-ATAC-seq protocol[7] with minor modifications. The cell pellet was resuspended in 100 µL cold Cell Lysis Buffer (10 mM Tris-HCl pH 7.4, 10 mM NaCl, 3 mM $MgCl_2$, 0.1% NP-40 alternative [Millipore Sigma]), spun as above, and the supernatant was discarded. Chromatin was then transposed by gentle resuspension of the nuclei pellet in Transposition Mix (25 µL 2X Tagment DNA Buffer, 16.5 µL 1x PBS, 0.1% Tween-20 [Promega], 0.01% Digitonin [Promega], 5 µL Tn5 Transposase [Illumina], 2.5 µL nuclease-free $H_2O$) and incubation at 30 min at 37°C with agitation at 1,000rpm. The reaction was terminated by the addition of 5 µL of 1% SDS (Promega). The transposed DNA was purified using the MinElute Reaction Cleanup Kit (Qiagen) with two sequential elutions into a final volume of 20 µL water, followed by 7-8 cycles of PCR amplification using PCR amplification mix (20µl Transposed DNA, 2.5 µL 25 µM Customized Nextera PCR Index Primer 1, 2.5 µL 25 µM Customized Nextera PCR Index Primer 2, 25 µL NEB High Fidelity 2x PCR Master Mix) under the following conditions (72°C for 5 minutes; 98°C for 30 seconds; 7-8 cycles of 98°C, 10 seconds, 63°C, 30 seconds, 72°C, 1 minute). Amplified DNA was then purified using a 1:1 ratio AMPure XP beads (Beckman Coulter) according to manufacturer specifications. The size and quality of each library were then evaluated by Bioanalyzer 2100 (Agilent) and quantified using qPCR (New England Biolabs). Libraries were sequenced in paired-end mode on the NextSeq 2000 platform to generate 2 x 50 bp reads using Illumina-supplied kits as appropriate. The sequence reads were processed using the ENCODE3 ATAC-seq pipeline (https://github.com/ENCODE-DCC/atac-seq-pipeline). Heatmaps and profiles were generated as described in the "ChIP-seq and data analysis" section.

**ChIP-seq library preparation**

We treated log-phase LOUCY$^{DH}$ and KOPT K1$^{DH}$ cells with DMSO or 500 nM dTAG$^{V}$-1 for 4 hours. Cells were harvested, centrifuged at 500g x 5 minutes at 4°C, washed with cold 1x PBS supplemented with 2 mM EDTA, and washed again with cold 1x PBS. Cell pellets were resuspended at 1e6 cells/mL in room temperature 1x PBS. Disuccinimidyl glutarate (DSG, ProteoChem) was added dropwise to the cell suspension at a final concentration of 2 mM, and samples were gently nutated for 30 minutes at room temperature. Formaldehyde (Millipore Sigma) was added to a final concentration of 1% with an additional 10 minutes of nutation. Crosslinking was quenched by the addition of glycine to a final concentration of 1 M and nutation for 5 minutes. Samples were centrifuged at 1200g x 5 minutes at 4°C and washed twice in cold 1x PBS with 0.5% BSA in Protein LoBind tubes (Eppendorf). Crosslinked cells were divided into 10e6 cell aliquots, and snap-frozen in liquid nitrogen and stored at -80°C for up to 1 year. Per ChIP, 10e6 crosslinked cells were lysed in 1 mL cold Cell Lysis Buffer (10 mM Tris-HCl pH 8.0, 10 mM NaCl, 0.2% NP-40 alternative [Millipore Sigma], 1 mM PMSF, 0.2x Protease Inhibitor Cocktail [Millipore Sigma] in $H_2O$) for 20 minutes with rotation at 4°C. Swollen cells were passed through a 27G needle 6 times to completely lyse cell membranes. Nuclei were pelleted by centrifugation at 2500g for 5 minutes at 4°C and resuspended in 0.33mL cold Nuclei Lysis Buffer (50 mM Tris-HCl pH 8.0, 10 mM EDTA, 1% SDS [Promega], 1 mM PMSF, 0.2x Protease Inhibitor Cocktail [Millipore Sigma] in $H_2O$) for 20 minutes with gentle agitation at 4°C. Samples were sonicated with the Bioruptor Pico (Diagenode) for 12-13 minutes (30 seconds on/off, “medium” mode). Sonication efficiency was checked by rapid decrosslinking of 1% input DNA (95°C x 15 minutes), cleanup with the ChIP Clean and Concentrator Kit (Zymo Research), and electrophoresis on a 2% agarose gel to ensure fragmentation to 200-500 bp. Sonicated chromatin was clarified by centrifugation at 15,000g x 10 minutes at 4°C in DNA LoBind tubes (Eppendorf). 1% of chromatin input was set aside and stored at -20°C for later use. The supernatant was diluted 10-fold with IP Dilution Buffer (20 mM Tris-HCl pH 8.0, 2 mM EDTA, 150 mM NaCl, 1% Triton X-100 [Roche], 0.01% SDS [Promega], 1 mM PMSF, 0.2x Protease Inhibitor Cocktail [Millipore Sigma] in $H_2O$). For each ChIP, 37.5 µL each of Protein A and Protein G Dynabeads (Invitrogen) were mixed together, washed twice with cold Blocking Solution (0.5% BSA, 0.1% Triton X-100 [Roche], 1 mM PMSF, 0.2x Protease Inhibitor Cocktail [Millipore Sigma] in 1x PBS), and conjugated with 10 µg of antibody in Blocking Solution for at least 2 hours at 4°C. All antibodies are reported in **Supplemental Table 14**.

Dynabead-antibody conjugates were washed twice again with cold Blocking Solution and added to the diluted chromatin, followed by end-over-end rotation at 4°C for at least 6 hours. After immunoprecipitation, bead-antibody-chromatin complexes were washed once with cold Wash Buffer 1 (20 mM Tris-HCl pH 8.0, 2 mM EDTA, 50 mM NaCl, 1% Triton X-100 [Roche], 0.1% SDS [Promega], 1 mM PMSF, 0.2x Protease Inhibitor Cocktail [Millipore Sigma] in $H_2O$), twice with cold High Salt Buffer (20 mM Tris-HCl pH 8.0, 2 mM EDTA, 500 mM NaCl, 1% Triton X-100 [Roche], 0.01% SDS [Promega], 1 mM PMSF, 0.2x Protease Inhibitor Cocktail [Millipore Sigma] in $H_2O$), once with cold LiCl Wash Buffer (10 mM Tris-HCl pH 8.0, 1 mM EDTA, 0.25M LiCl, 1% NP-40 alternative [Millipore Sigma], 1% Sodium Deoxycholate, 1 mM PMSF, 0.2x Protease Inhibitor Cocktail [Millipore Sigma] in $H_2O$), and twice with cold 1x TE Buffer (Promega). Chromatin was eluted in 100 µL of Elution Buffer (100 mM $NaHCO_3$, 1% SDS [Promega] in $H_2O$) for 15 minutes at 65°C with agitation at 1,000rpm. Two sequential elutions were pooled for a final volume of 200 µL. Stored 1% chromatin inputs were diluted to 200 µL with Elution Buffer. 300 mM NaCl and 100 µg/mL RNase A (Millipore Sigma) were added to eluted chromatin and inputs and incubated at 60°C for 1-2 hours. 300 µg/mL Proteinase K (Invitrogen) was then added and incubated at 65°C to complete decrosslinking for 12-16 hours. DNA was purified using the ChIP Clean and Concentrator Kit (Zymo Research) per manufacturer specifications. ChIP-qPCR was performed to verify target enrichment using Power Sybr Green PCR Master Mix (Applied Biosystems) on either Viia 7 or QuantStudio 5 platforms. ChIP-seq libraries were prepared using the NEBNext Ultra II DNA Library Prep Kit (New England Biolabs) with NEBNext Multiplex Oligos for Illumina (New England Biolabs). Libraries were purified and concentrated using AMPure XP beads (Beckman Coulter) according to manufacturer specifications. Libraries were quantified by Qubit (Invitrogen), and their size and quality were evaluated by Bioanalyzer 2100 (Agilent). Libraries were sequenced in paired-end mode on an Illumina Novaseq X Plus platform at Novogene (2 x 150 bp reads) or on an Illumina NextSeq 2000 platform (2 x 50 bp reads).

**TT-seq library preparation**

Log-phase LOUCY$^{DH}$ and KOPT K1$^{DH}$ cells (15e6 cells per condition) were treated with DMSO or 500 nM dTAG$^{V}$-1 for 4 hours and labeled with 500µM 4-thiouridine (4SU) (MedChemExpress) for 5 minutes. Cells were centrifuged at 500g x 5 minutes at 4°C and immediately resuspended in 3 mL TRIzol Reagent (Invitrogen). Total RNA was extracted per manufacturer specifications using MaXtract tubes (Qiagen) to assist with separation of the aqueous layer. 500 ng of 4SU-labeled Drosophila Schneider 2 cells total RNA spike-in was mixed with 100µg of purified 4SU-labeled KOPT K1 or LOUCY total RNA. Mixed RNA was fragmented using a final concentration of 0.2M NaOH for 18 minutes and neutralized by adding Tris-HCl (pH 6.8) to 0.5M. Fragmented

RNA was purified by isopropanol precipitation and biotinylated in 300µL of Biotinylation Mix (10 mM HEPES pH 7.5, 1 mM EDTA, 10 µg MTSEA-biotin [Biotium] in DEPC-treated $H_2O$) for 1 hour at room temperature and purified with phenol/chloroform/isoamyl alcohol (25:24:1 [v:v:v]) extraction. Biotinylated RNA was then denatured at 65°C x 10 minutes, followed by rapid cooling on ice for 5 minutes. The denatured biotinylated RNA was bound to Dynabeads MyOne Streptavidin C1 (Invitrogen) at room temperature for 30 minutes in 1x Binding and Wash Buffer (100 mM Tris-HCl pH 7.5, 1 M NaCl, 10 mM EDTA, 0.05% Tween-20 in DEPC-treated $H_2O$). Supernatant was removed and bead-RNA complexes were washed 4 times with 1x Binding and Wash before elution with 1x Binding and Wash Buffer supplemented with 100 mM DTT. Eluted RNA was purified by isopropanol precipitation. Quality of total RNA extraction, RNA fragmentation (aiming for RNA size distribution between 25 and 500 bp), and purified 4SU-labeled RNA were determined using Agilent TapeStation RNA ScreenTape (Agilent). Strand-specific sequencing libraries were generated with the Illumina Stranded Total RNA Prep (Illumina) and IDT for Illumina RNA UD Indexes Set A, Ligation (Illumina). Library size was determined using Agilent TapeStation High Sensitivity DNA ScreenTape (Agilent). Libraries were pooled and sequenced on the Illumina NextSeq 2000 platform to generate 2 x 50 bp reads using Illumina sequencing reagents according to the manufacturer specifications.

**Micro-C library preparation**

We treated log-phase LOUCY$^{DH}$ and KOPT K1$^{DH}$ cells (10e6 cells per biological replicate) with DMSO or 500 nM dTAG$^{V}$-1 for 4 hours. Cells were harvested, centrifuged at 500g x 5 minutes at 4°C, washed twice with cold 1x PBS, and resuspended in room temperature 1x PBS at 1e6 cells/mL. Formaldehyde (Millipore Sigma) was added dropwise at a final concentration of 1%

followed by gentle nutation for 10 minutes at room temperature. The reaction was quenched with 0.375M Tris-HCl pH 7.5 and nutation for 5 minutes. Cells were centrifuged at 1,000g x 5 minutes, washed twice with 1x PBS, and then resuspended in room temperature 1x PBS at 1e6 cells/mL. DSG (ProteoChem) was then added dropwise at a final concentration of 3 mM followed by gentle nutation for 40 minutes at room temperature. The reaction was quenched with 0.375M Tris-HCl pH 7.5 and nutation for 5 minutes. Cells were centrifuged at 1,000g x 5 minutes, split into 2 technical replicates (5e6 cells each), washed twice with cold 1x PBS with 0.5% BSA in Protein LoBind tubes (Eppendorf), snap-frozen and stored at -80°C. For all biological replicates, 2 technical replicates were processed in parallel. Each 5e6 cell technical replicate was resuspended in 250 µL cold complete MNase Buffer 1 (10 mM Tris-HCl pH 7.5, 50 mM NaCl, 5 mM $MgCl_2$, 1 mM $CaCl_2$ in $H_2O$ supplemented with 0.2% NP-40 alternative [Millipore Sigma] and 1x EDTA-free cOmplete protease inhibitor cocktail [Roche]) and chilled on ice for 20 minutes prior to centrifugation at 2,000g x 10 minutes at 4°C. Pellet was resuspended in complete MNase Buffer 1 with 15 U MNase (Worthington Biochemical) to digest chromatin for 20-35 minutes at 37°C to attain 9:1 mononucleosome:dinucleosome fragments. The duration of MNase digestion was tested for each biological replicate based on a time course experiment. Digestion was halted with 4 mM EGTA pH 10 and incubation for 10 minutes at 65°C. Samples were centrifuged and washed 3 times in 250 µL of cold complete MNase Buffer 2 (10 mM Tris-HCl pH 7.5, 50 mM NaCl, 10 mM $MgCl_2$ in $H_2O$ supplemented with 100 µg/mL recombinant BSA [New England Biolabs]). Pellets were resuspended in 100 µL 1x NEBuffer 2.1 (New England Biolabs); 10 µL was set aside as an input control to test ligation efficiency. The remaining 90 µL reaction solution was dephosphorylated by adding 10 U r-SAP (New England Biolabs), mixed at 37°C for 45 minutes, and inactivated at 65°C for 5 minutes. For end-chewing of dephosphorylated DNA, we pooled two technical replicates and added 5 µL 10x NEBuffer 2.1, 5 µL 100 mM ATP (Thermo Scientific), 7.5 µL 100 mM DTT, 11.5 µL $H_2O$, and 50 U T4 PNK (New England Biolabs), mixed at 37°C for 15 minutes, added 80 U Large Klenow Fragment (New England Biolabs), and mixed for an additional 15 minutes at 37°C (final volume 250 µL). DNA ends were labeled by adding the following to the reaction solution: 50 µM each Biotin-dATP and Biotin-dCTP (Jena Bioscience), 50 µM each of dTTP and dGTP (Promega), 0.5x T4 DNA Ligase Buffer (New England Biolabs), 20 µg recombinant BSA (New England Biolabs), and $H_2O$ to a final volume of 400 µL. The reaction solution was intermittently mixed (shake 1 minute/rest 3 minutes) at 25°C for 45 minutes and then inactivated by adding 30 mM EDTA and heating to 65°C for 20 minutes. To increase complexity of ligated fragments, reactions were then split into 2 tubes, centrifuged, and washed twice with 1x T4 DNA Ligase Buffer. Each pellet was then resuspended in the Proximity Ligation Solution (1x T4 DNA Ligase Buffer, 50 µg recombinant BSA, 10,000 U T4 DNA Ligase [New England Biolabs], 1x EDTA-free cOmplete protease inhibitor cocktail, and $H_2O$ to 500 µL). Samples were then nutated in 0.65 mL LoBind tubes (Eppendorf) overnight (~16 hours) at room temperature. Samples were centrifuged for 10 minutes at 4°C, resuspended in Exonuclease Solution (1x NEBuffer 1 [New England Biolabs], 300 U Exonuclease III [New England Biolabs], and $H_2O$ to 146 µL), and mixed at 37°C for 15 minutes to remove unligated ends. Ligated DNA and input controls underwent decrosslinking by adding 1% SDS (Promega), 300 mM NaCl, 100 µg/mL RNase A (Millipore Sigma), and 300 µg/mL Proteinase K (Invitrogen), followed by incubation at 65°C for 12-16 hours. Ligated DNA fragments were purified by phenol:chloroform:isoamyl alcohol (25:24:1) extraction and ethanol precipitation. Proximity ligation efficiency was analyzed by electrophoresis on a 2% agarose gel to ensure increased dinucleosome abundance compared with input controls. The two technical replicates were pooled, cleaned twice with 0.9x AMPureXP beads (Beckman Coulter), and eluted into 150 µL 10 mM Tris-HCl. We next prepared 5 µL Dynabeads MyOne Streptavidin C1 (Invitrogen) per pooled sample (1 biological replicate) by washing twice in 1x TBW Buffer (5 mM Tris-HCl pH 7.5, 0.5 mM EDTA, 1 M NaCl, 0.05% Tween-20 in $H_2O$) and resuspension in 150 µL 2x BW Buffer (10 mM Tris-HCl pH 7.5, 1 mM EDTA, 2M NaCl in $H_2O$). Beads were mixed with eluted DNA, rotated end-over-end at room temperature for 30 minutes prior to magnetic separation, washed twice with 1x TBW Buffer at 55°C, and washed once with 100 µL 10 mM Tris-HCl. Bead-DNA complexes were then resuspended into 50 µL 10 mM Tris-HCl, and Micro-C libraries were prepared using the NEBNext Ultra II DNA Library Prep Kit (New England Biolabs) with NEBNext Multiplex Oligos for Illumina (New England Biolabs) using on-bead PCR amplification. PCR cycles were determined by a "side" PCR reaction to maintain linear amplification of libraries. Size selection of dinucleosomes was then performed by using 0.55x AMPure XP beads to remove large DNA fragments and then mixing the supernatant with 0.7x AMPure XP beads to remove small DNA fragments. Size distribution and quality of libraries was analyzed using Bioanalyzer 2100 (Agilent Technologies). Shallow sequencing libraries were prepared in paired-end mode on the NextSeq 2000 platform to generate 2 x 50 bp reads. Deep sequencing of high-quality libraries was then performed in paired-end mode on the Illumina Novaseq X Plus platform at Novogene to generate 2 x 150 bp reads.

**Micro-C compartment analysis**

We used cooltools v0.7.1 to compute cis eigenvector values from 100 kb binned matrices from DMSO and dTAG$^{V}$-1 treatments. These data were used to generate saddleplots reflecting the average observed/expected contact frequency between AA, BB, AB, and BA interactions, as well as saddle strength profiles reflecting the strength of compartmentalization.

**Micro-C domain analysis**

Domains were called using rGMAP[8] v1.4 with 10 kb-binned contact matrices. Domain calls were merged between DMSO and dTAG$^{V}$-1 treatments, deduplicated, filtered to remove domains with start/end coordinates within 80 kb and/or domains <100 kb. We defined boundaries as 120 kb windows flanking start/end coordinates of each domain and calculated insulation scores at boundaries using cooltools with a 120 kb sliding window. We then compared the minimum insulation scores at all boundaries for DMSO and dTAG$^{V}$-1 treatments.

**Loop calling and quantification**

We used cooltools.dots to call loops using merged contact maps for DMSO or dTAG$^{V}$-1 treatments in either the LOUCY$^{DH}$ or KOPT K1$^{DH}$ cell lines. Loops were separately called using 10 kb, 5 kb, and 2 kb contact maps for each cell line and treatment using the following parameters: max_loci_separation=2_000_000, clustering_radius=20_000, lambda_bin_fdr=0.05 and n_lambda_bins=50. We used "rounded donuts" and "lowleft" kernels to identify loops and merged loop calls from DMSO and dTAG$^{V}$-1 for each resolution and each cell line. DMSO-dTAG$^{V}$-1 merged loop lists were then merged across all resolutions for each cell line, with the smallest resolution coordinates retained for redundant loops, generating distinct merged lists of loop calls for LOUCY$^{DH}$ or KOPT K1$^{DH}$ cells. Loop strength was then quantified for DMSO and dTAG$^{V}$-1 treatments within each cell line by calculating the observed/locally adjusted expected value. Loop strength was calculated for each loop using the resolution at which the loop was identified, and loops with strengths of 0, NA, or infinite were removed. This resulted in a final consensus list of 38,752 loops in LOUCY$^{DH}$ and 30,724 loops in KOPT K1$^{DH}$. We then calculated the $\log_2$(fold change [FC]) for each loop such that negative values represented weakened loops and positive values represented strengthened loops upon dTAG$^{V}$-1 treatment. A $\log_2$(FC) cutoff of >|0.5| was used to define strengthened or weakened loops.

### Supplemental Figures and Legends

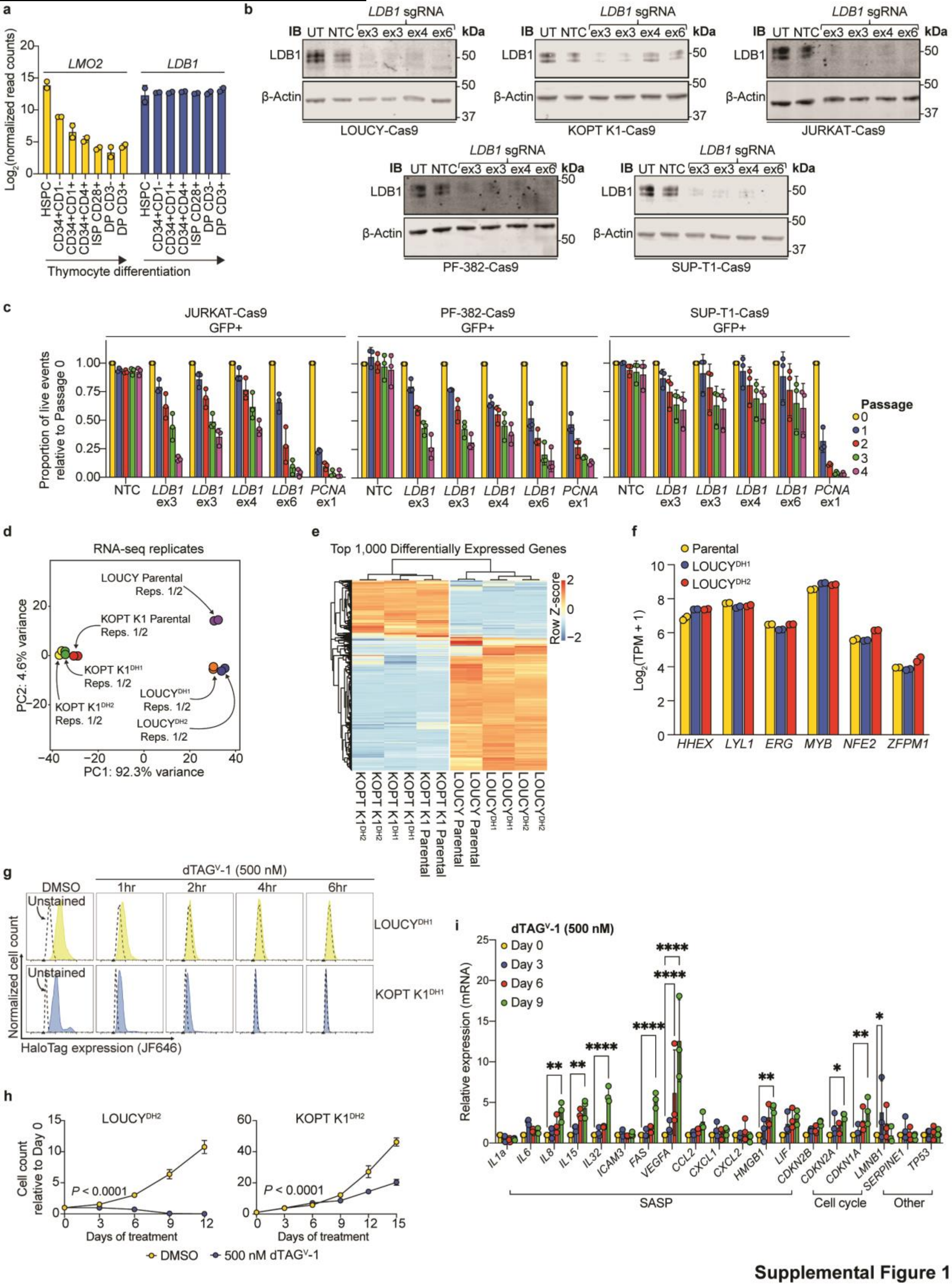


**Supplemental Figure 1**

**Supplemental Fig. S1. Orthogonal validation of LDB1 knockout and dTAG systems. a,** DESeq2-normalized read counts for *LMO2* and *LDB1* in normal T-cell precursors based on postnatal thymocyte RNA-seq data[52]. **b,** Representative Western blots depict LDB1 and β-actin protein expression in Cas9-expressing T-ALL cell lines transduced with control or *LDB1*-targeting sgRNAs. **c,** Proportion of GFP+ JURKAT-, PF-382- or SUP-T1-Cas9 cells after transduction with NTC, *PCNA*-targeting (positive control) or *LDB1*-targeting sgRNAs; values normalized to Passage 0 for each sgRNA. **d,** Principal component analysis of RNA-seq from LOUCY and KOPT K1 parental cells and dTAG-HaloTag subclones; biological replicates are denoted by arrows. **e,** Heatmap with unsupervised hierarchical clustering of LOUCY and KOPT K1 parental cells and dTAG-HaloTag subclones based on the top 1,000 differentially expressed genes; color scale, row Z-score. **f,** Transcripts per million (TPM) for known LDB1 target gene transcripts in LOUCY parental cells and dTAG-HaloTag subclones. **g,** Time-dependent changes to HaloTag signal detection in $\text{LOUCY}^{\text{DH1}}$ and KOPT $\text{K1}^{\text{DH1}}$ cells upon $\text{dTAG}^{\text{V}}$-1 treatment using the J646 Janelia Fluor HaloTag Ligand. **h,** $\text{LOUCY}^{\text{DH2}}$ and KOPT $\text{K1}^{\text{DH2}}$ cell counts upon serial DMSO or $\text{dTAG}^{\text{V}}$-1 treatment (500 nM); cell counts normalized to Day 0; *P*-values from two-way ANOVA with post hoc Tukey test. **i,** Relative mRNA expression of senescence-associated genes shown from RT-qPCR in KOPT $\text{K1}^{\text{DH1}}$ cells serially treated with $\text{dTAG}^{\text{V}}$-1; *P*-values from one-way ANOVA with post hoc Dunnett's test compared with Day 0 (only *P*-values for Day 9 are shown). $N = 3$ biological replicates for all experiments, except for RNA-seq ($N = 2$ each). HSPC, hematopoietic stem and progenitor cell; ISP, immature single-positive T-cell; DP, double-positive thymocyte; UT, untransduced; NTC, non-targeting control; ex, exon; IB, immunoblot; P, passage; PC, principal component; J646, Janelia Fluor 646; SASP, senescence-associated secretory phenotype.

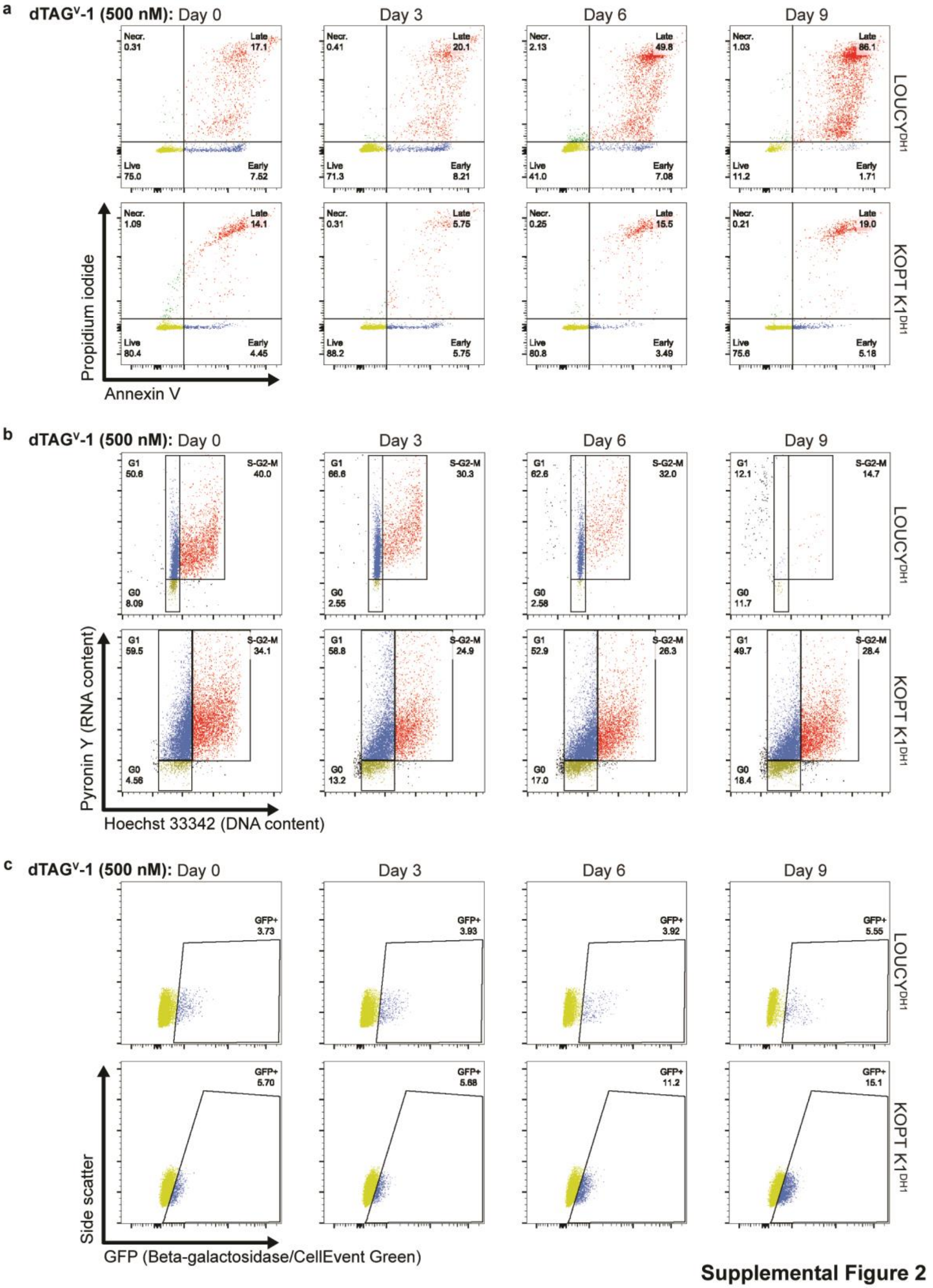


**Supplemental Fig. S2. Representative flow cytometry scatterplots. a,** Scatter plots depict

Annexin V versus propidium iodide staining in $LOUCY^{DH1}$ or KOPT $K1^{DH1}$ cells to measure early/late apoptosis and necrosis upon serial $dTAG^{V}$-1 treatment. Gating schema applies to **Fig. 1h**. **b,** Scatter plots depict Hoechst 33342 versus pyronin Y staining for DNA and RNA content, respectively in $LOUCY^{DH1}$ or KOPT $K1^{DH1}$ cells to measure cell cycle distribution upon serial $dTAG^{V}$-1 treatment. Gating schema applies to **Fig. 1g**. **c,** Scatter plots depict GFP (β-galactosidase/CellEvent Green) versus side scatter in $LOUCY^{DH1}$ or KOPT $K1^{DH1}$ cells to measure β-galactosidase expression as a marker of senescence upon serial $dTAG^{V}$-1 treatment. Gating schema applies to **Fig. 1i**.

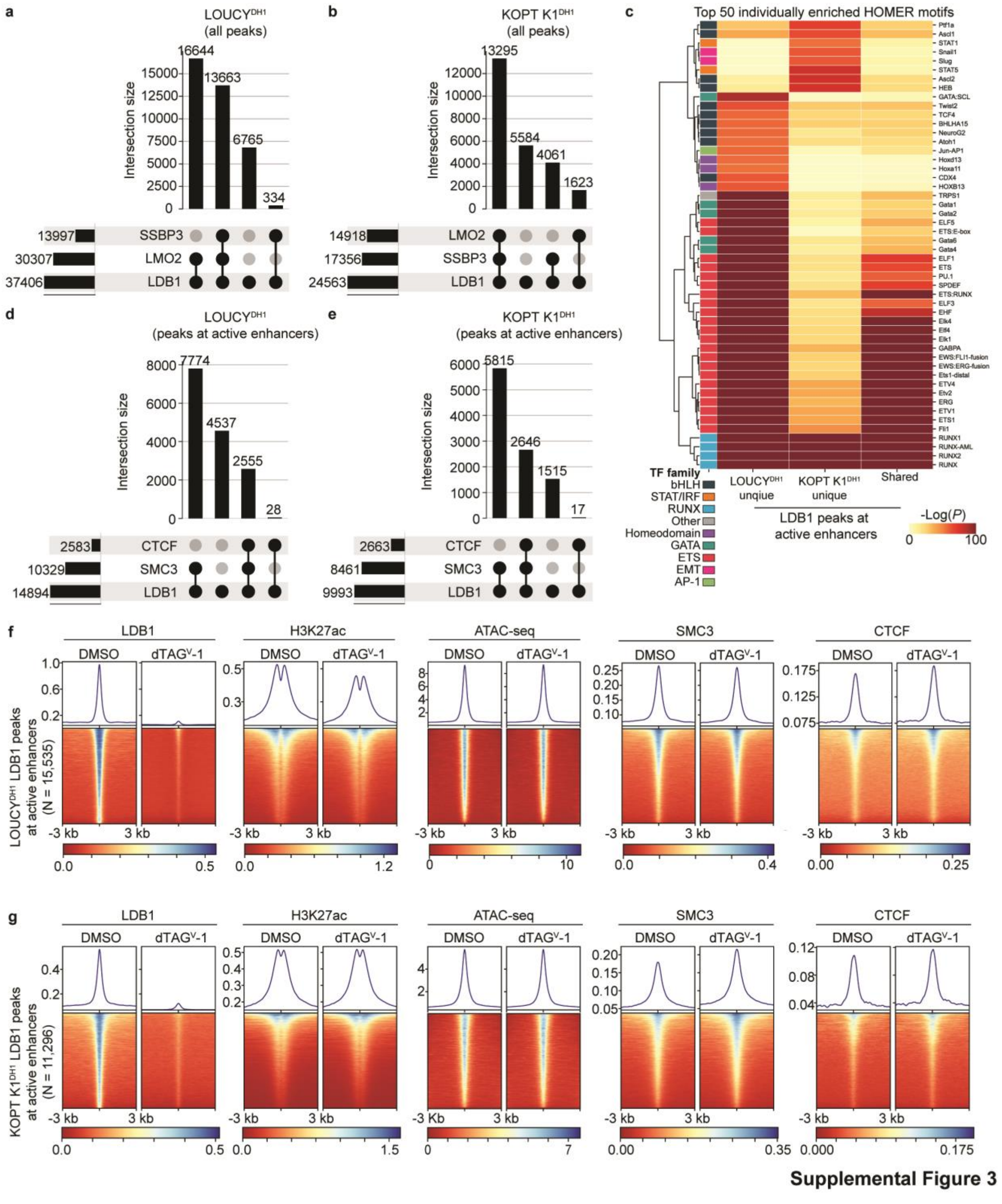


**Supplemental Fig. S3. Subtype-specific transcription factors govern LDB1/LMO2 complex recruitment to enhancers. a,** UpSet plot of LMO2 and SSBP3 ChIP-seq peak overlap with LDB1 peaks in LOUCY$^{DH1}$. **b**, Similar to **a**, but depicting KOPT K1$^{DH1}$. **c,** Heatmap of HOMER Known

Motif enrichment at LOUCY$^{DH1}$-specific, KOPT K1$^{DH1}$-specific or shared LDB1 peaks at enhancers categorized by transcription factor (TF) family; individual motif calls are labeled. **d,** UpSet plot of CTCF and SMC3 ChIP-seq peak overlap with LDB1 peaks at active enhancers in LOUCY$^{DH1}$. **e**, Similar to **d**, but depicting KOPT K1$^{DH1}$. **f,** Heatmaps of LOUCY$^{DH1}$ ChIP-seq and ATAC-seq signal upon acute LDB1 depletion; profiles centered over all LDB1 peaks at enhancers from DMSO-treated cells. **g**, Similar to **f,** but depicting KOPT K1$^{DH1}$ ChIP-seq and ATAC-seq data. Biological replicates per cell line and treatment: LDB1 ChIP-seq, $N = 4$; LMO2 ChIP-seq, $N = 2$; SSBP3 ChIP-seq, $N = 2$; H3K27ac ChIP-seq, $N = 2$; ATAC-seq, $N = 3$.

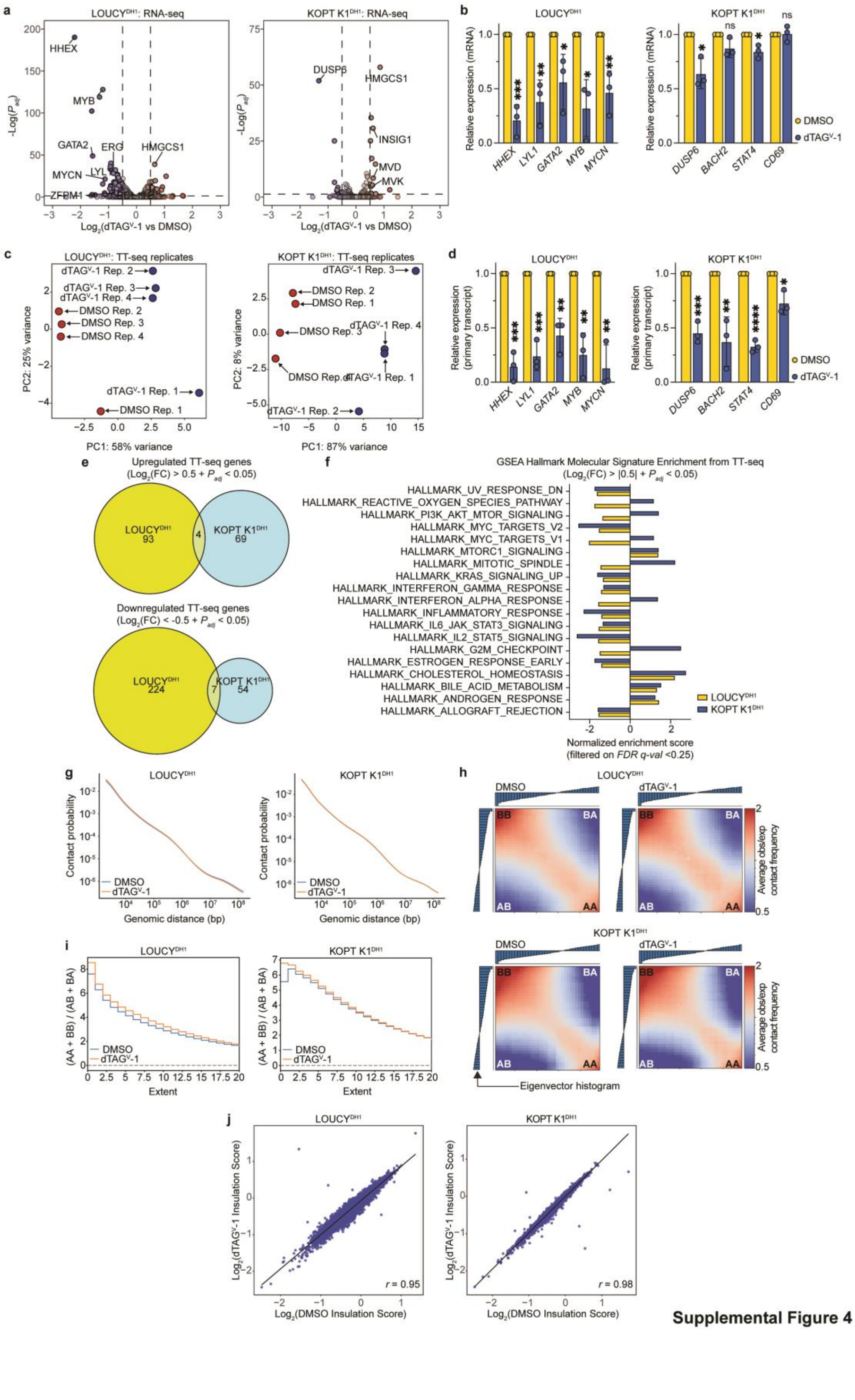


**Supplemental Figure 4**

**Supplemental Fig. S4. Acute LDB1 depletion disrupts T-ALL subtype-specific oncogenic transcriptional programs but minimally impacts larger scale chromatin structure. a,** Volcano plots of gene expression changes measured by RNA-seq upon acute LDB1 depletion; relevant genes are denoted (blue, downregulated; red, upregulated). **b,** Relative mRNA expression of LDB1-dependent genes shown from RT-qPCR in $LOUCY^{DH1}$ and KOPT $K1^{DH1}$ upon acute LDB1 depletion; *q*-values from unpaired Student's *t*-test with Benjamini-Hochberg correction. **c,** Principal component analysis comparing TT-seq replicates from $LOUCY^{DH1}$ and KOPT $K1^{DH1}$. **d,** Relative primary transcript expression of LDB1-dependent genes shown from RT-qPCR in $LOUCY^{DH1}$ and KOPT $K1^{DH1}$ upon acute LDB1 depletion; *q*-values from unpaired Student's *t*-test with Benjamini-Hochberg correction. **e,** Venn diagrams showing overlap between $LOUCY^{DH1}$ and KOPT $K1^{DH1}$ for TT-seq upregulated and downregulated TT-seq gene sets. **f,** Summary of GSEA based on TT-seq data from $LOUCY^{DH1}$ and KOPT $K1^{DH1}$ upon acute LDB1 depletion. **g,** Contact decay curves from $LOUCY^{DH1}$ and KOPT $K1^{DH1}$ Micro-C upon acute LDB1 depletion. **h,** Saddle plots with overlayed Eigenvector histograms from $LOUCY^{DH1}$ and KOPT $K1^{DH1}$ Micro-C upon acute LDB1 depletion (A-A, A-B, B-B and B-A interactions are labeled). **i,** Saddle strength profiles from $LOUCY^{DH1}$ and KOPT $K1^{DH1}$ Micro-C upon acute LDB1 depletion. **j,** Scatter plots of $log_2$-transformed topologically associating domain (TAD) boundary insulation scores from $LOUCY^{DH1}$ and KOPT $K1^{DH1}$ Micro-C upon acute LDB1 depletion; Pearson correlation coefficient (*r*) shown. $N = 2$ biological replicates for RNA-seq; $N = 3$ biological replicates for RT-qPCR; $N = 4$ biological replicates for TT-seq per cell line and treatment; $N = 3$ pooled biological replicates for Micro-C per cell line and treatment. PC, principal component; FC, fold change; GSEA, gene set enrichment analysis; FDR, false discovery rate; A, active chromatin compartment; B, inactive chromatin compartment.

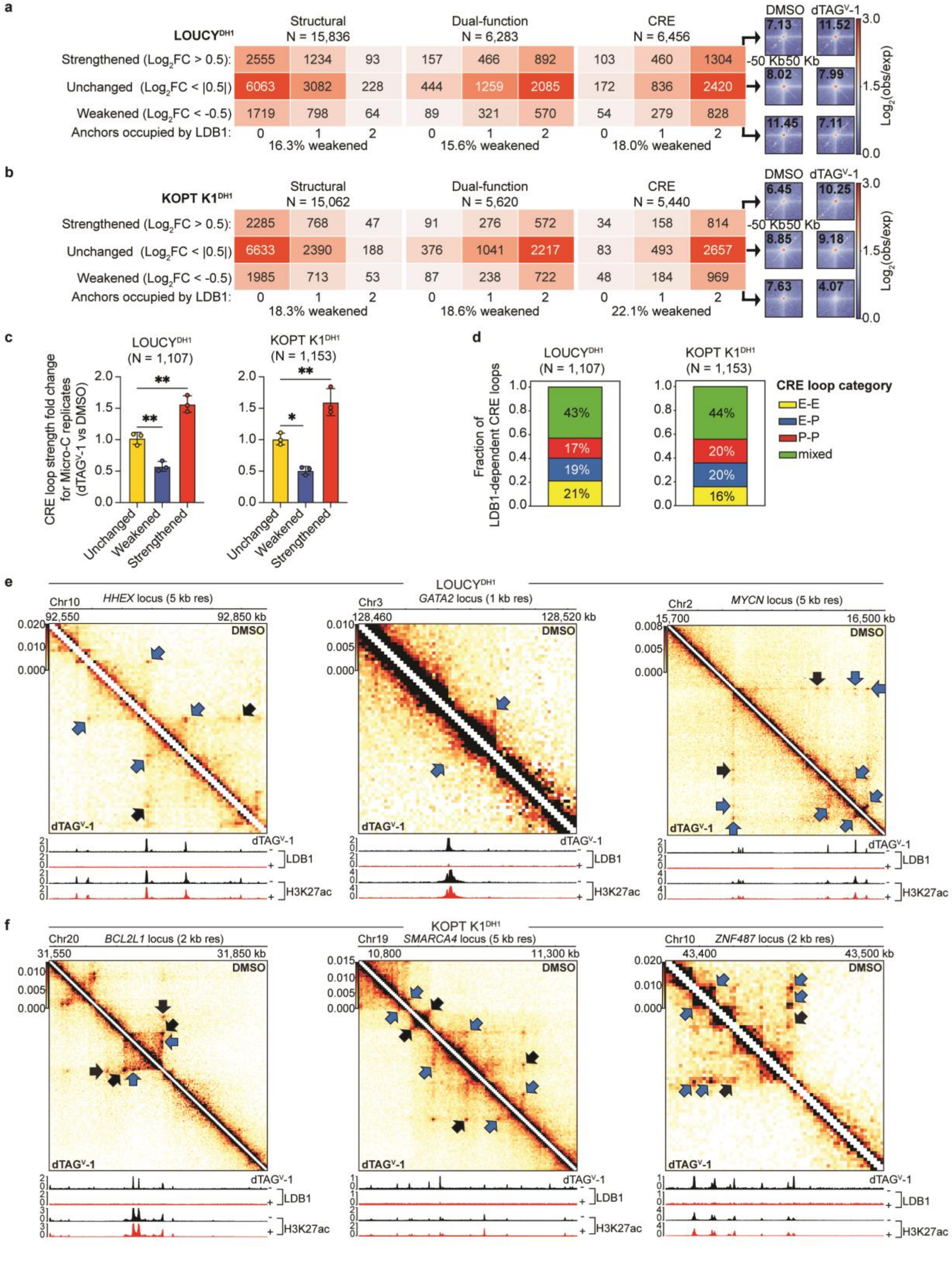


**Supplemental Figure 5**

**Supplemental Fig. S5. Acute LDB1 depletion preferentially disrupts cis-regulatory element loops. a,** Numbers of structural, dual-function and CRE loops in LOUCY$^{DH1}$ that are weakened, unchanged or strengthened upon acute LDB1 depletion, stratified by LDB1 presence at loop anchors; 2 kb resolution pileup plots shown for CRE loops. **b,** Similar to **a,** but depicting loop changes for KOPT K1$^{DH1}$. **c,** Fold change in observed/expected CRE loop strength for individual Micro-C replicates (each dot represents one biological replicate); *P*-values from one-way ANOVA with post hoc Tukey test. **d,** Distribution of LDB1-dependent CRE loops by enhancer (E), promoter (P) or mixed connectivity. **e,** Representative Micro-C contact maps from LOUCY$^{DH1}$, depicting CRE loop changes upon 4-hour DMSO and dTAG$^{V}$-1 treatments (blue arrow, weakened LDB1-dependent loops; black arrow, unchanged/strengthened LDB1-independent loops), with aligned LDB1 and H3K27ac ChIP-seq tracks (black, DMSO; red, dTAG$^{V}$-1). **f**, Similar to **e,** but depicting Micro-C contact maps from KOPT K1$^{DH1}$. $N$ = 3 pooled biological replicates per cell line and treatment, except panel **c**, which depicts individual biological replicates. CRE, cis-regulatory element.

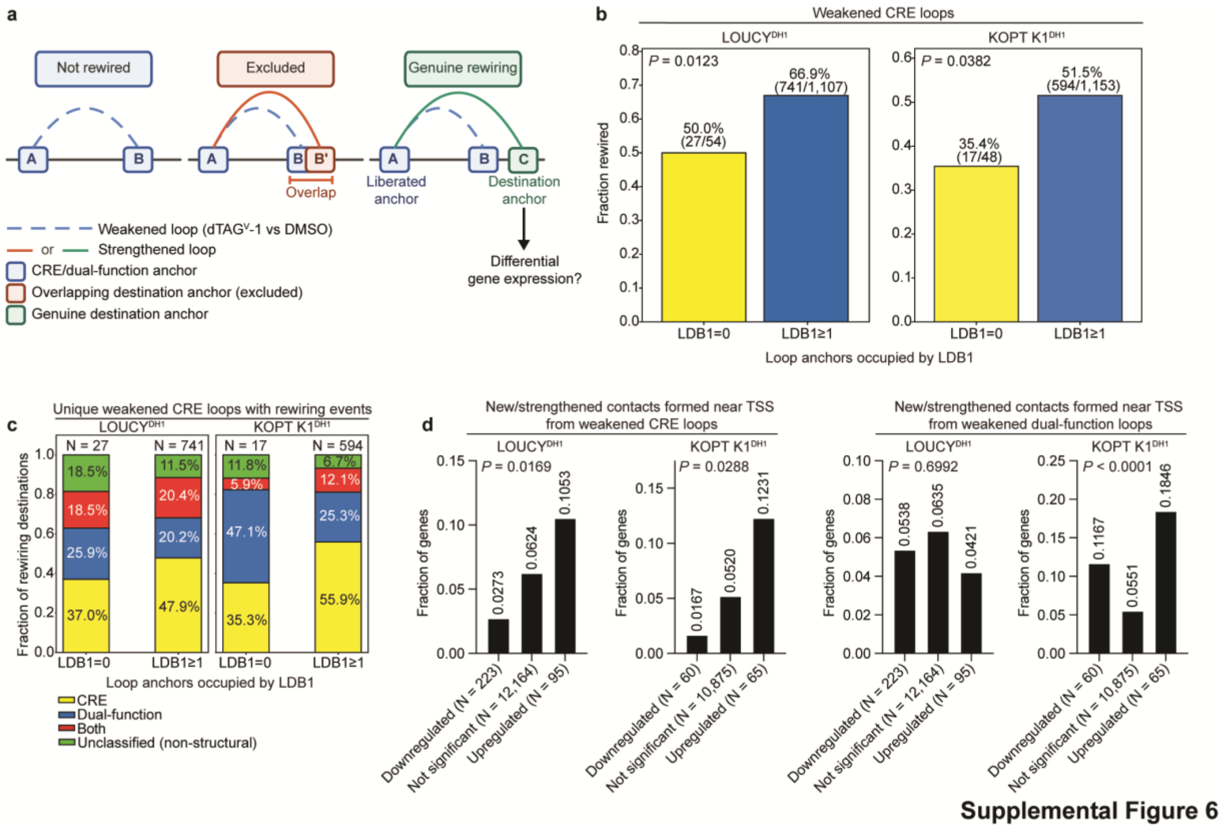


**Supplemental Figure S6. Acute LDB1 depletion promotes rewiring of cis-regulatory element loops. a,** Schematic of CRE loop rewiring analysis (anchor A, liberated anchor; anchor B, lost/weakened contact/anchor; anchor C, destination/gained contact/anchor). **b,** Proportion of weakened CRE loops that share an anchor with a distinct strengthened loop (rewiring) upon acute LDB1 depletion; *P*-values from Fisher's exact test; **c,** Classification of rewiring destinations from weakened CRE loops. **d**, Fraction of TT-seq differentially expressed genes overlapping with a rewired loop; *P*-value from Fisher's exact test. CRE, cis-regulatory element; TSS, transcription start site.

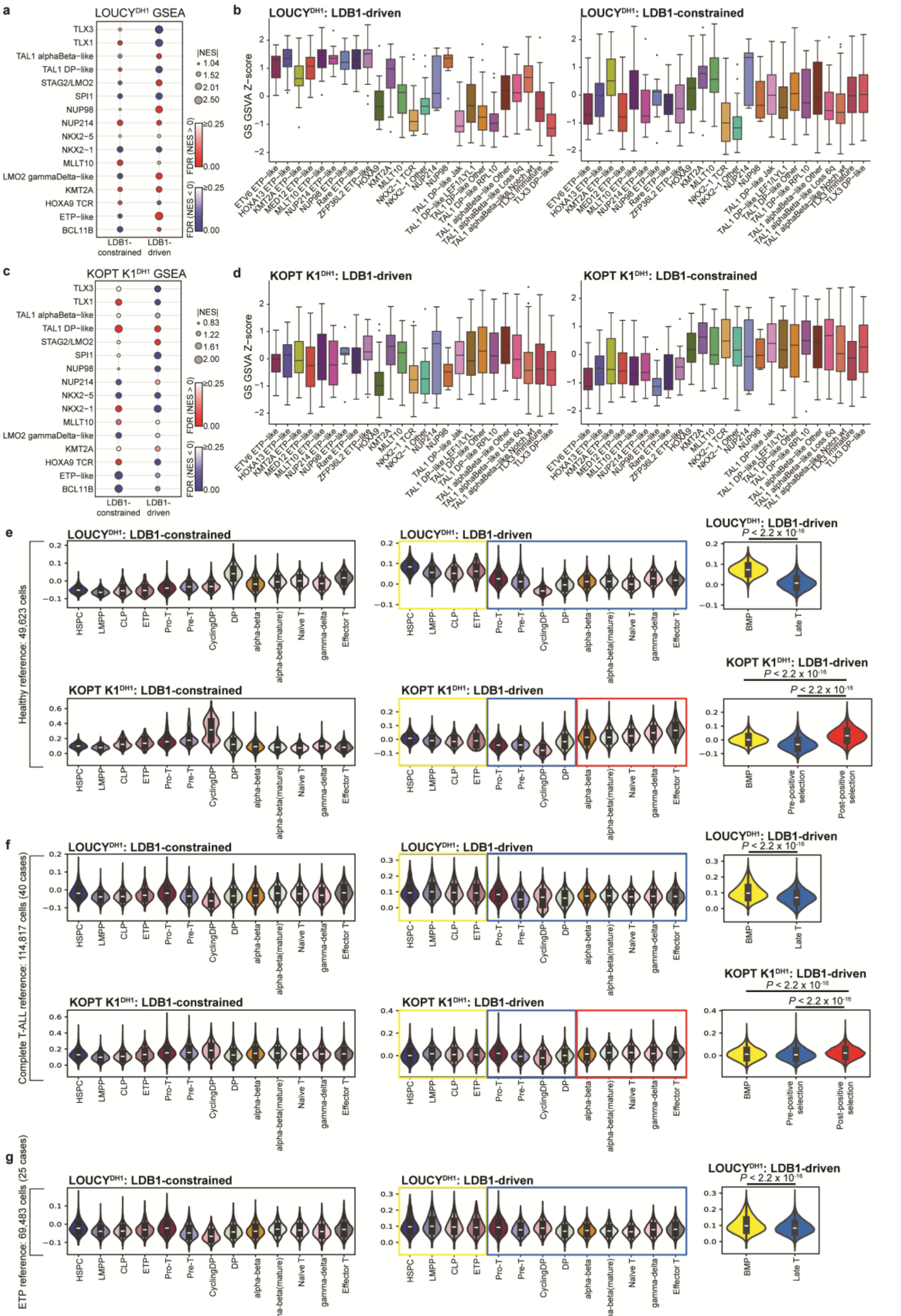


**Supplemental Figure 7**

**Supplemental Figure S7. LDB1 drives T-ALL subtype identities and constrains against alternative transcriptional states. a,** Summary of GSEA for LDB1-constrained and LDB1-driven gene sets from $\text{LOUCY}^{\text{DH1}}$ TT-seq in primary T-ALL whole transcriptome data (*N* = 1,335 patients), stratified by molecular subtype. **b,** Box-and-whisker plots depict GSVA enrichment-like scores of LDB1-constrained and LDB1-driven gene sets from $\text{LOUCY}^{\text{DH1}}$ TT-seq in primary T-ALL data, stratified by driver genetic lesion. **c,** Similar to **a**, but depicting KOPT $\text{K1}^{\text{DH1}}$ GSEA. **d,** Similar to **b**, but depicting KOPT $\text{K1}^{\text{DH1}}$ GSVA. **e,** Violin plots depict normalized expression of TT-seq-derived LDB1-driven gene sets in a healthy pediatric hematopoiesis scRNA-seq reference dataset (*N* = 49,623 cells), classified as bone marrow progenitor-like (BMP-like) versus late T-cell or BMP-like versus pre- or post-positive selection T-cells (outline colors correspond with violin shading in summary plot); *P*-value from Mann-Whitney test. **f**, Similar to **e**, but depicting normalized expression in a T-ALL scRNA-seq reference dataset (*N* = 114,817 cells). **g**, Similar to **e, f,** but depicting normalized expression of $\text{LOUCY}^{\text{DH1}}$ LDB1-driven gene sets in an ETP ALL scRNA-seq reference dataset (*N* = 69,483 cells). GSEA, gene set enrichment analysis; GSVA, gene set variation analysis; DP, double-positive thymocyte; ETP, early thymic precursor.

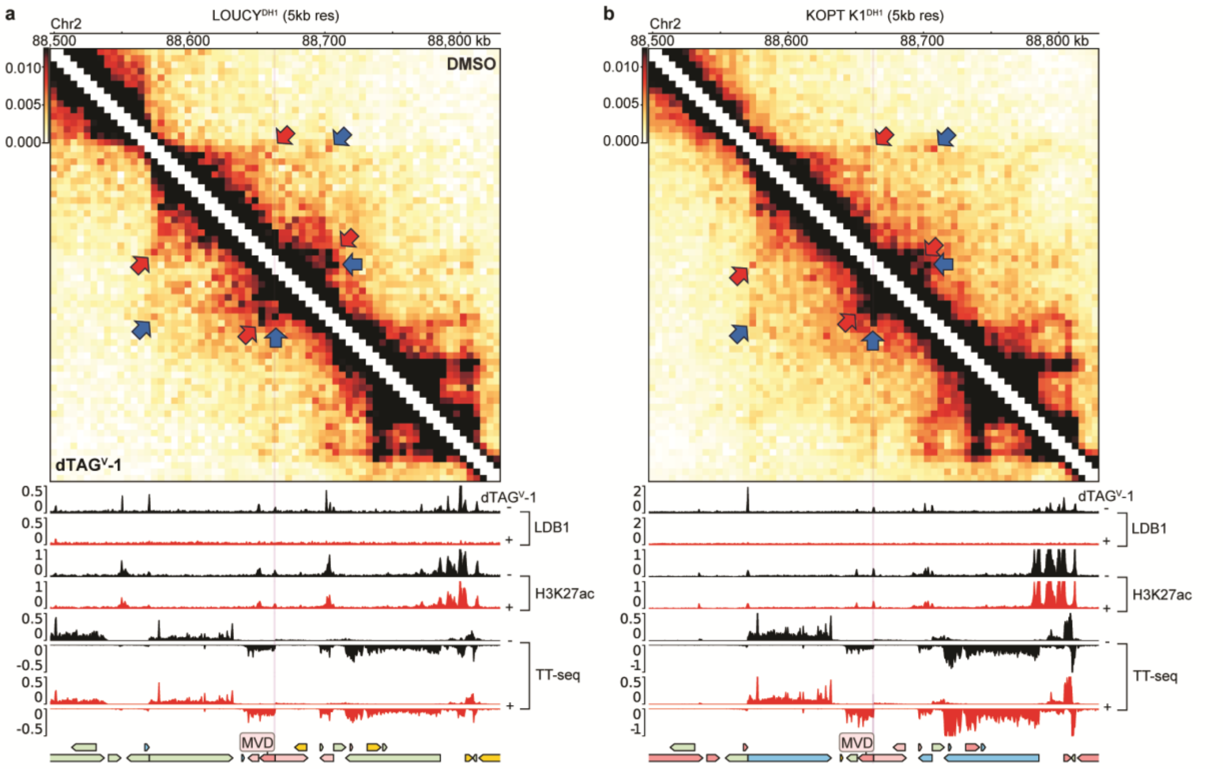


**Supplemental Figure 8**

**Supplemental Fig. S8. Representative Micro-C contact maps for mevalonate pathway gene chromatin loop rewiring. a,** 5 kb resolution Micro-C contact map of the *MVD* locus in LOUCY$^{DH1}$, depicting CRE loop rewiring events upon 4-hour DMSO and dTAG$^{V}$-1 treatments (blue arrow, weakened loops; red arrow, strengthened loops; purple shading, *MVD* promoter), with aligned LDB1 and H3K27ac ChIP-seq and TT-seq tracks (black, DMSO; red, dTAG$^{V}$-1). **b,** Similar to **a,** but depicting the *MVD* locus in KOPT K1$^{DH1}$ at 5 kb resolution.

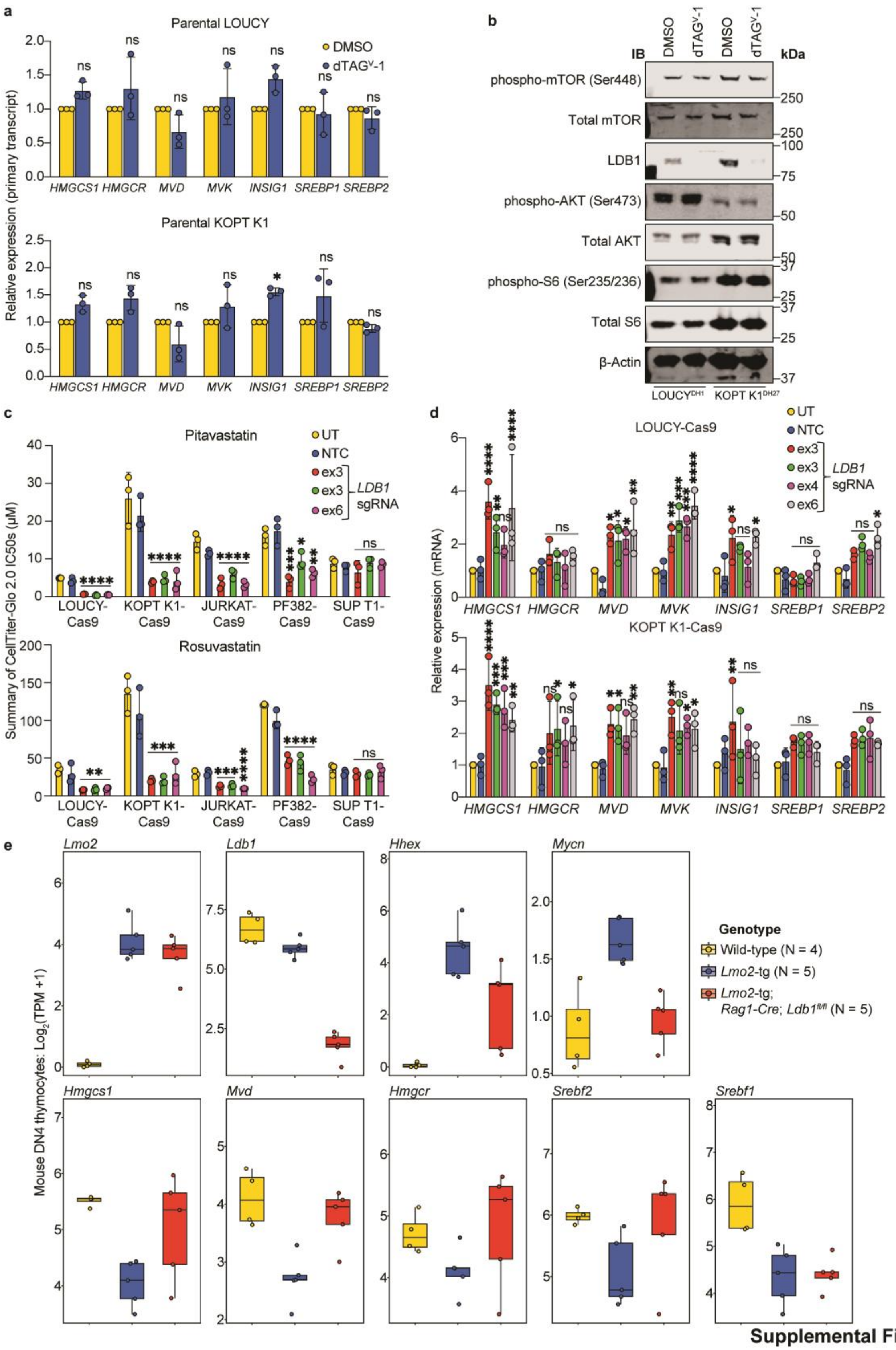


**Supplemental Figure 9**

**Supplemental Figure S9. Mevalonate pathway phenotypic changes induced by LDB1 loss are conserved in human T-ALL cell lines and mouse models of T-ALL. a,** Relative primary transcript expression of mevalonate pathway genes shown from RT-qPCR in parental LOUCY and KOPT K1 cells treated upon 4-hour treatment with DMSO or dTAG$^{V}$-1; *q*-values from unpaired Student's *t*-test with Benjamini-Hochberg correction. **b,** Representative Western blot depicts LDB1; β-Actin; and phospho- and total mTOR, AKT and S6 protein expression upon acute LDB1 depletion. **c,** Summary IC50 values from CellTiter-Glo 2.0 assays for Cas9-expressing T-ALL cell lines treated with statins concomitant with *LDB1* knockout; *P*-values from one-way ANOVA with post hoc Dunnett's test compared with untransduced cells. **d,** Relative mRNA expression of mevalonate pathway genes shown from RT-qPCR in LOUCY- and KOPT K1-Cas9 cells transduced with control or *LDB1*-targeting sgRNAs; *P*-values from one-way ANOVA with post hoc Dunnett's test compared with untransduced cells. **e,** Box-and-whisker plots depicting *Lmo2*, *Ldb1,* known target oncogene and key mevalonate pathway gene expression ($log_2$[TPM+1]) from RNA-seq[38] in mouse DN4 thymocytes with *Lmo2* overexpression (*Lmo2*-tg, $N$ = 5), *Lmo2* overexpression with conditional LDB1 knockout (*Lmo2*-tg; *Rag1-Cre*; *Ldb1$^{fl/fl}$*, $N$ = 5) or wild-type *Lmo2/Ldb1* status (wild-type, $N$ = 4). $N = 3$ biological replicates for all other experiments. NTC, non-targeting control; ex, Exon; IB, immunoblot; IC50, half-maximal inhibitory concentration; DN4, double-negative 4 thymocytes.
